# Hi-TrAC reveals fine-scale chromatin structures organized by transcription factors

**DOI:** 10.1101/2022.06.01.494329

**Authors:** Shuai Liu, Yaqiang Cao, Kairong Cui, Qingsong Tang, Keji Zhao

## Abstract

The three-dimensional genomic structure plays a critical role in gene expression, cellular differentiation, and pathological conditions. It is pivotal to elucidate fine-scale chromatin architectures, especially interactions of regulatory elements, to understand the temporospatial regulation of gene expression. In this study, we report Hi-TrAC as a proximity ligation free, robust, and sensitive technique to profile genome-wide chromatin interactions at high-resolution among accessible regulatory elements. Hi-TrAC detects chromatin looping among accessible chromatin regions at single nucleosome resolution. With almost half-million identified loops, we constructed a comprehensive interaction network of regulatory elements across the genome. After integrating chromatin binding profiles of transcription factors, we discovered that cohesin complex and CTCF are responsible for organizing long-range chromatin loops, related to domain formation; whereas ZNF143 and HCFC1 are involved in structuring short-range chromatin loops between regulatory elements, which directly regulate gene expression. Thus, we developed a new methodology to identify a delicate and comprehensive network of cis-regulatory elements, revealing the complexity and a division of labor of TFs in chromatin looping for genome organization and gene expression.

## Introduction

The genome is organized into higher-order chromatin structures^1–6^. Each chromosome occupies a discrete territory in the nucleus^7^. Based on the spatial separation of active and inactive phases, the chromatin is partitioned into A and B compartments respectively^7–9^. Self-associating chromatin assembles into ∼1 Mb sized topologically associating domains (TADs)^10–12^, which contain nested sub-TADs with the size of several hundred kb^13, 14^. Chromatin domains are assembled by chromatin interaction loops, which are organized by CTCF and cohesin complex through a loop-extrusion process^15–26^. Interactions between transcriptional regulatory elements are important for orchestrating gene expression^27–43^. Knowledge of the detailed enhancer-promoter interaction network is important for understanding the fine-tuning of cell activities^44–46^. A sensitive and efficient technique is highly desired for elucidating genome-wide fine structures, particularly enhancer-promoter interactions. It is generally accepted that cohesin complex catalyzes chromatin folding into loops anchored by CTCF binding^18–23, 43, 47–50^, whereas several other chromatin factors, including ZNF143 and YY1, have been shown to facilitate chromatin loop formation^8, 51–58^. However, it is not clear how these architectural proteins orchestrate chromatin looping at different scales of genome organization. In this study, we profiled chromatin interactions among accessible regions using a new technique termed as Hi-TrAC (h<u>i</u>ghly sensitive <u>tr</u>ansposase-mediated <u>a</u>nalysis of <u>c</u>hromatin) and elucidated the activities of CTCF, RAD21, HCFC1, and ZNF143 in long-range versus short-range chromatin looping.

## Results

### Technical improvements of Hi-TrAC

Hi-TrAC originated from the Trac-looping (Transposase-mediated analysis of chromatin looping) method with substantial improvements^59^. Hi-TrAC takes advantage of DNA transposase Tn5’s ability of utilizing a specially designed bivalent linker to covalently bridge spatially proximal open chromatin regions, thus eliminating chromatin fragmentation and proximity ligation steps required for 3C (chromosome conformation capture)-based techniques (**Fig. 1a, Methods**). We also designed a strategy to eliminate the rolling cycle amplification and dilute ligation in large volumes in Trac-looping, enabling us to reduce the starting material of 100 million cells to as few as 10 thousand cells and shorten library construction time from 7 days to 2 days.

**Fig. 1.**
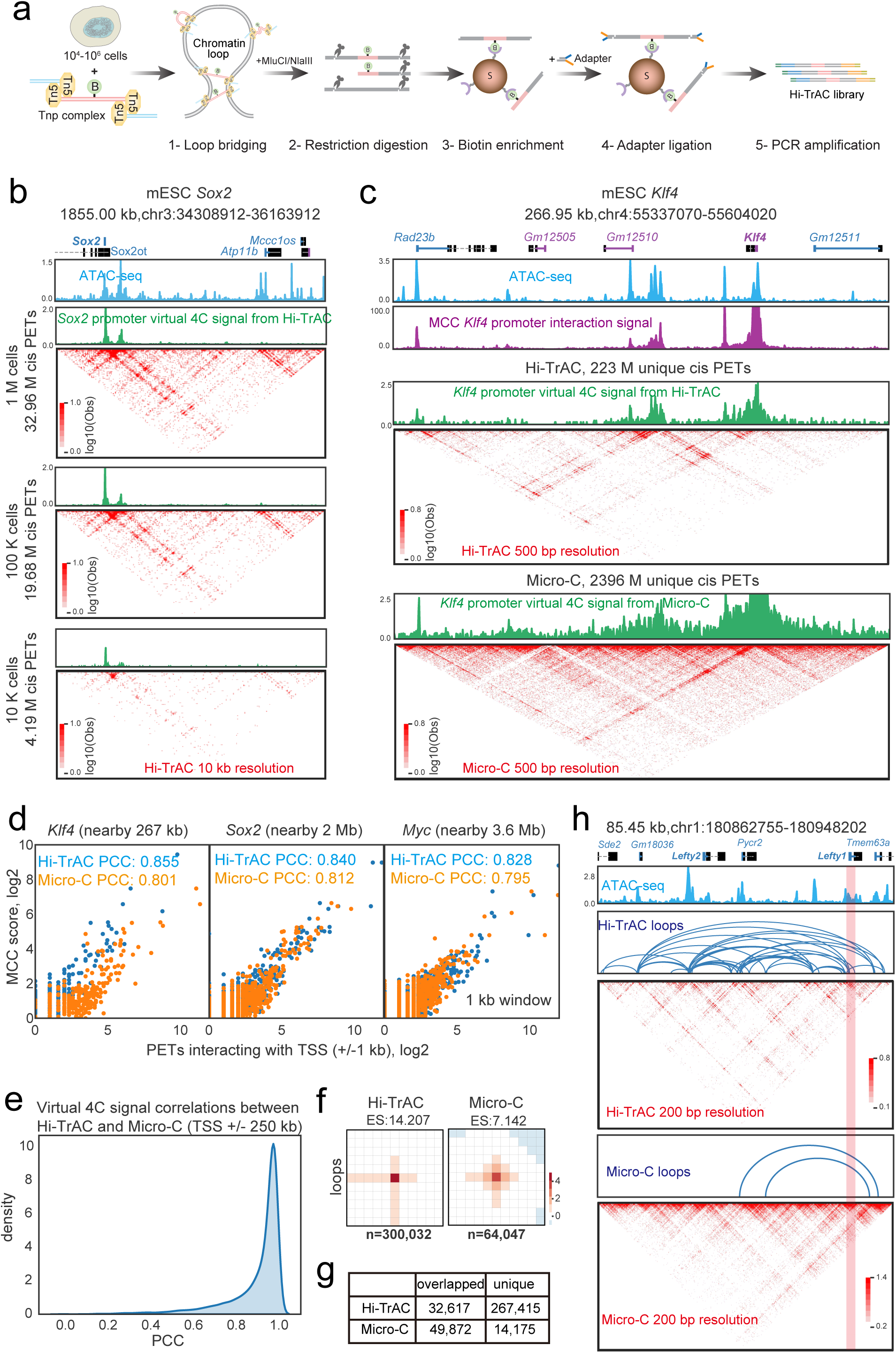
Mapping genome-wide regulatory interactions at high resolution by Hi-TrAC. **(a)** Experimental scheme of Hi-TrAC. Following bridging chromatin loops using the Tnp-biotinylated bivalent ME linker complex in formaldehyde fixed cells, the DNA is cleaved with restriction enzymes MluC I and Nla III. The bridged genomic regions are enriched using streptavidin beads and PCR-amplified for sequencing after ligation of a universal adaptor. **(b)** Hi-TrAC reproducibly detects interactions around the *Sox2* gene locus from 10^6^, 10^5^, and 10^4^ E14 mESCs. ATAC-seq data were obtained from GSM1830114^114^. Hi-TrAC virtual 4C signals were generated by only keeping PETs interacting with the +/-1kb TSS region of *Sox2* gene and displayed as the piled up 1D signal. Interacting PETs were shown as dots below the 4C-like signals. The genomic annotations are shown on the top of the panel. The visualization was performed with cLoops2 plot module. **(c)** Comparison of interactions around *Klf4* gene locus from pooled MCC^62^, Hi-TrAC, and Micro-C^64^ data in mESCs. Only intra-chromosomal PETs from Hi-TrAC and Micro-C were used for comparisons. **(d)** Correlation analysis between interactions detected by MCC with the virtual 4C signals from Hi-TrAC or Micro-C data around *Klf4*, *Sox2*, and *Myc* loci, with the viewpoint set as the +/-1kb of TSS. **(e)** Distribution of correlations between the virtual 4C signals from Hi-TrAC and Micro-C around the promoter regions of all protein-coding genes. The promoter defined as a region +/-1Kb upstream and downstream of a TSS was set as the viewpoint. Only PETs with any end located in the promoter region were kept, and a region of 250Kb upstream and downstream of TSSs was set as the comparing region. PCC stands for Pearson Correlation Coefficient. **(f)** Aggregation analysis of Hi-TrAC and Micro-C loops. Hi-TrAC loops were called by the cLoops2 callLoops module and requiring at least 20 PETs (**Supplemental Table 2**), and Micro-C loops were called by HiCCUPS. **(g)** Overlaps of Hi-TrAC and Micro-C loops. **(h)** Genome Browser snapshot of the *lefty* locus, showing the distribution of ATAC-seq peaks and chromatin loops detected by Hi-TrAC and Micro-C as well as the interaction matrices at a 200 bp resolution.

### Hi-TrAC outperforms other techniques at detecting chromatin loops

We first benchmarked Hi-TrAC with the most advanced Hi-C variants Micro-C and Micro-Capture-C (MCC), which produce high-resolution chromatin structure maps^60–62^. Starting with 0.01, 0.1, and 1 million mouse embryonic stem cells (mESCs) (**Supplemental Table 1**), as exemplified by the *Sox2* gene locus, Hi-TrAC reproducibly detected similar chromatin interaction profiles (**Fig. 1b**). To compare with MCC data, we extracted all paired-end tags ( PETs) linking to the promoters of *Klf4*, *Sox2,* and *Myc* genes from Hi-TrAC and Micro-C data and displayed them as virtual 4C signals. Hi-TrAC detected chromatin interactions at these promoters were highly comparable to those detected by MCC and Micro-C, from both visual inspection and a quantitative correlation analysis (**Fig. 1c-d**). Moreover, the virtual 4C signals from Hi-TrAC and Micro-C data were highly correlated at all gene promoters in the genome (**Fig. 1e**). With a higher signal-to-noise ratio, Hi-TrAC even showed higher correlation with MCC than Micro-C, and detected more details of fine chromatin structures (**Fig. 1c-d and Extended Data Fig. 1a).** These results indicate that Hi-TrAC is a robust and reliable technique for detecting interactions among chromatin regulatory regions.

Hi-TrAC performed a little more sensitively than Micro-C to detect interactions at different genomic distances (**Extended Data Fig. 1b**). We identified 300k significant chromatin loops from 223 million unique intra-chromosomal PETs (cis-PETs) in Hi-TrAC, compared to 64k loops from 2,396 million unique cis-PETs in Micro-C (**Supplemental Table 2**). The aggregation analysis showed that Hi-TrAC loops displayed a higher enrichment score (ES) than Micro-C loops, indicating a higher signal-to-noise ratio in Hi-TrAC data (**Fig 1f**). About 80% of Micro-C loops were identified by Hi-TrAC, whereas only 11% of Hi-TrAC loops were covered by Micro-C (**Fig. 1g and Extended Data Fig. 1c**). As exemplified by the looping profiles at loci of functionally important genes, including *Myc* (**Extended Data Fig. 1a**), *Lefty* (**Fig. 1h),** and *Nanog* **(Extended Data Fig. 1d**), more significant loops were identified by Hi-TrAC owing to its high signal-to-noise ratio. Especially at the promoters of *Lefty1* and *Nanog* genes, specific loops could only be identified by Hi-TrAC, but missed by Micro-C (**Fig. 1h and Extended Data Fig. 1d**). Generally, Micro-C unique loops are distal weaker loops (**Extended Data Fig. 1e-f**). The majority of both Micro-C (76%) and Hi-TrAC (92%) loop anchors were enriched at accessible chromatin regions as characterized by ATAC-seq peaks (**Extended Data Fig. 1g**) The rest did not show features of repressive chromatin (**Extended Data Fig. 1h**), suggesting that significant chromatin loops detected by these techniques were mainly between accessible regulatory elements. Together, these results indicate that Hi-TrAC is a sensitive method for detecting chromatin loops among active regulatory regions.

To further evaluate the performance of Hi-TrAC in elucidating chromatin structures, we applied Hi-TrAC to GM12878 cells, a human cell-line whose genome architecture had been extensively studied by various techniques (**Supplemental Table 1**). To obtain a comprehensive interaction map, we pooled the data from all experimental replicates, resulting in 822 million raw reads and 117 million unique intra-chromosomal PETs (**Supplemental Table 1**). As shown in the two-dimension (2D) heatmap at different resolutions, we compared the genome architecture map generated by Hi-TrAC with available maps built by in situ Hi-C^8^, CTCF^32^ and RAD21 ChIA-PET^51, 63^, capture Hi-C^64^, H3K27ac HiChIP^65^ and cohesin HiChIP^66^ (**Extended Data Fig. 2, some data are not shown**). With relatively low sequencing depth, chromatin domain-like structures and loops could be clearly detected by Hi-TrAC at different resolutions; especially for identifying significant chromatin loops, Hi-TrAC data had a much higher signal-to-noise ratio than other methods; even at 200 bp resolution, the fine architectural details of super-enhancers could also be observed, which were not clear in the maps generated by other techniques (**Extended Data Fig. 2**).

To systematically compare the genome-wide highest resolution that these techniques can achieve, we calculated the coverage of PETs with different bin sizes. With the threshold of more than 50% of the PETs not being singleton PETs, only H3K27ac HiChIP could achieve a similar resolution to Hi-TrAC (**Extended Data Fig. 3).** We then performed sub-samplings of Hi-TrAC data to estimate the required sequencing depth for reaching a desired resolution. The analysis indicated that with 60 million intra-chromosomal PETs, about 300 million raw reads, genome-wide 1 kb resolution could be achieved (**Extended Data Fig. 4a**), and fine-scale architectures could also be identified at 200 bp resolution at a subset of genomic regions including super-enhancers (**Extended Data Fig. 4b-c**).

### Hi-TrAC detects comprehensive interaction network of cis-regulatory elements

To explore how the fine-scale chromatin architectures are organized for individual cis-regulatory elements, we further analyzed the chromatin looping revealed by Hi-TrAC data. In GM12878 cells, as exemplified by the *SPI1* locus, which encodes the key lymphoid cell development-related ETS family transcription factor PU.1,we observed typical dot-to-dot chromatin loops pattern revealed by Hi-TrAC, formed between the distal and proximal regulatory elements (**Fig. 2a)**. Globally, with at least 10 PETs supporting a loop, we called 91,042 high-confident loops in GM12878 cells (**Fig. 2b and Supplemental Table 3**), much more than other techniques (**Extended Data Fig. 5a**).

**Fig. 2.**
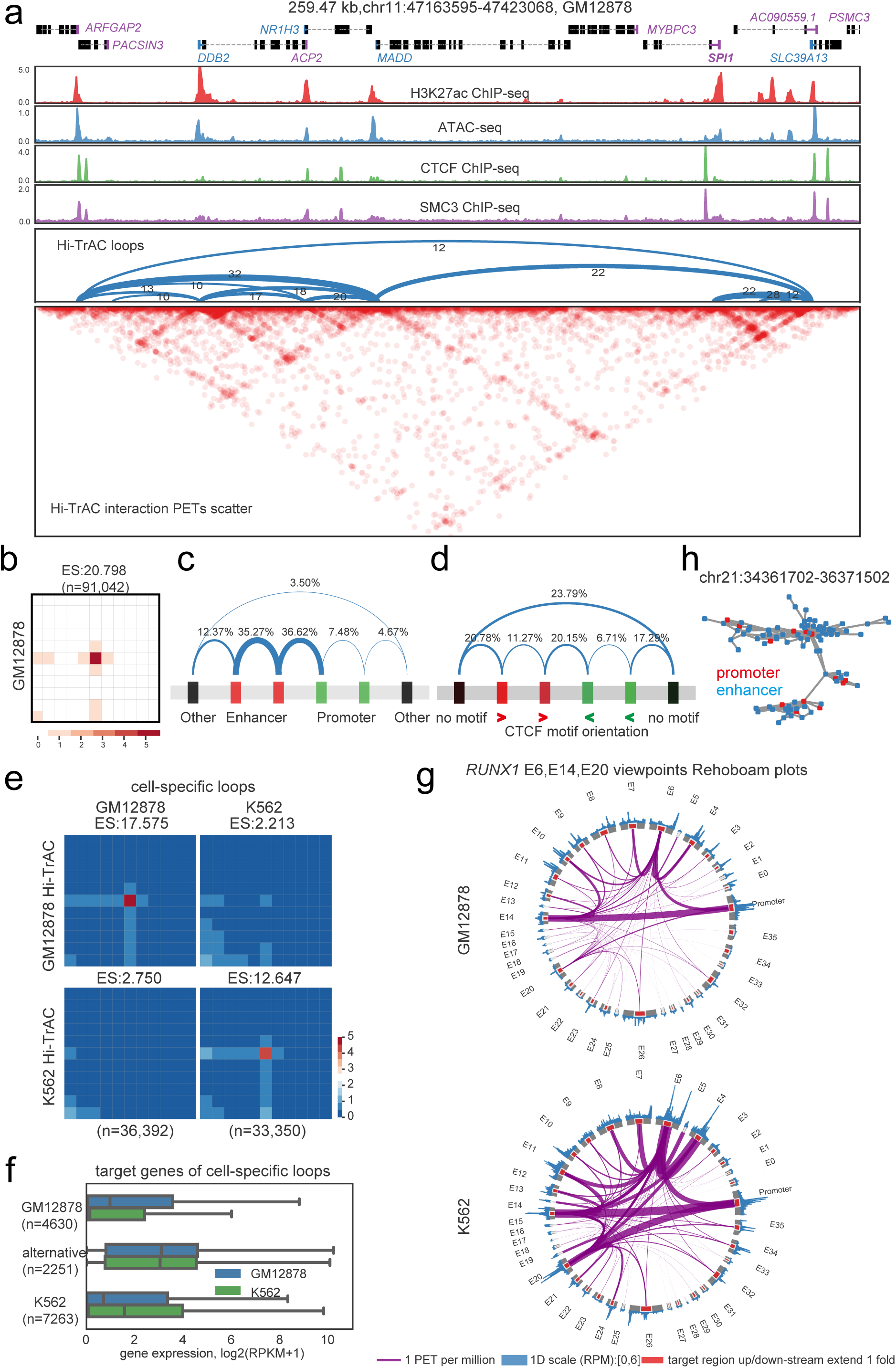
Chromatin looping networks constructed with Hi-TrAC data. **(a)** Genome Browser snapshot showing the chromatin loops detected by Hi-TrAC around the *SPI1* gene in GM12878 cells. Loops are shown as arches, and the numbers of PETs are also shown for each loop. Loops were called by the cLoops2 callLoops module, requiring at least ten supportive PETs. The interaction matrices are displayed at the bottom. **(b)** Aggregation analysis of 91,042 loops called from Hi-TrAC data in GM12878 (**Supplemental Table 3**). Interacting PETs in loops and their nearby regions (5-folds upstream and downstream of loop anchors) as matrices were averaged as the aggregated heatmap. ES stands for enrichment score, indicating the interaction signal enrichment compared to neighbor regions. The analysis was performed with cLoops2 agg module. **(c)** Summary of categories of GM12878 Hi-TrAC loops with regard to putative cis-regulatory elements, including enhancers, promoters and others. **(d)** Summary of categories of GM12878 Hi-TrAC loops with regard to the orientation of CTCF motifs at the two anchors of a loop. **(e)** Aggregation analysis of cell-specific loops in GM12878 and K562 (**Supplemental Table 4**). Differentially enriched loops were called with the cLoops2 callDiffLoops module. **(f)** The distribution of expression levels of the genes associated with cell-specific loops, and the genes with promoters looping with alternative enhancers between GM12878 and K562. The numbers of genes for each category were indicated. **(g)** Rehoboam plots showing the differences in promoter-enhancer interactions at *RUNX1* gene locus in GM12878 and K562 cells. Interactions from the viewpoints of enhancers E6, E14 and E20 are shown. **(h)** An example of the longest connected sub-network consisting of enhancers and promoters on Chromosome 21 in GM12878. Enhancers and promoters form complex connections as nature scale-free regulatory networks.

We compared various features of Hi-TrAC loops with those detected by other methods. Generally, Hi-TrAC loops were shorter in the distance, with loop anchors concentrated at promoters and enhancers (**Extended Data Fig. 5b-e)**. The majority of them were enhancer-promoter and enhancer-enhancer loops (**Fig. 2c**). Compared to other datasets, the CTCF motif orientations at Hi-TrAC loop anchors were more diverse (**Extended Data Fig. 5f and Fig. 2d**), with a small fraction of them in convergent orientation, and these loops appeared to be more distant (**Extended Data Fig. 5g**), suggesting these loops were likely to be related with domain formation. Many Hi-TrAC loop anchors did not have CTCF motifs (**Fig. 2d and Extended Data Fig. 5f**), suggesting different functions and organization mechanisms of these loops, and Hi-TrAC captured more versatile loops.

To investigate the relationship between chromatin loops and cell identity and activity, we further performed Hi-TrAC in another human cell-line, K562 cells (**Supplemental Table 1**). In total, 98,850 chromatin loops were identified in K562 cells (**Supplemental Table 3**). We hypothesized that cell-specific chromatin loops may control cell-specific gene expression. To test this, we compared the looping profiles in GM12878 and K562 cells, and identified 36,392 and 33,350 cell type-specific loops, respectively (**Supplemental Table 4**). Cell-specific loops showed significant interaction signal enrichment in corresponding cells (**Fig. 2e**). We identified 4,630 genes associated with GM12878-specific promoter loops and 7,263 genes associated with K562-specific promoter loops. Meanwhile, we also identified 2,251 genes that displayed alternative promoter-enhancer looping between GM12878 and K562 cells. Genes associated with GM12878-specific loops expressed at higher levels in GM12878 cells than in K562 cells; and vice versa (**Fig. 2f**). However, there were no significant differences in expression for the genes displaying alternative looping between GM12878 and K562 cells (**Fig. 2f**). We validated Hi-TrAC identified cell-specific loops with datasets from other methods through aggregation analysis, including in situ Hi-C, H3K27ac HiChIP and RAD21 ChIA-PET (**Extended Data Fig. 6**), demonstrating that Hi-TrAC successfully detected cell activity related differential chromatin loops.

The comprehensive regulatory interaction network was exemplified by several representative gene loci. EBF1 is a key transcription factor in B lymphopoiesis, which is expressed specifically in GM12878 cells, whereas GATA1 is a master transcription factor in erythropoiesis that is only expressed in K562 cells. *EBF1* and *GATA1* gene loci exhibited unique interaction patterns in corresponding cells, consistent with their expression profiles (**Extended Data Fig. 7a-b**). A previous elegant CRISPRi screening study identified multiple regulatory elements for *GATA1* expression^67^. Interestingly, taking the *GATA1* promoter as the viewpoint, the Hi-TrAC virtual 4C signals correlated well with the CRISPRi score (**Extended Data Fig. 7c-d**), indicating the robustness of detecting functionally relevant regulatory interactions by Hi-TrAC.

Take another K562 specific key transcription factor gene *RUNX1* as an example (**Extended Data Fig. 7e**). Although the comprehensive regulatory network can be presented with interaction contact matrix heatmaps and loop arc plots, it is too complicated for visual inspection for each individual *cis*-regulatory element. Thus, we designed a Rehoboam plot to visualize chromatin loops of a specific genomic region, which clearly revealed individual loops connecting different cis-regulatory elements: the interactions were much stronger between the promoter and enhancers E6 and E20 in K562 than in GM12878 cells, which thus might be responsible for its expression in K562 cells (**Fig. 2g**).

We integrated all the enhancer and promoter loops, and generated a comprehensive regulatory interaction network (**Fig. 2h**). The degree of connection of enhancers and promoters in the network fitted the scale-free network power-law, revealing the complexity of the regulatory network (**Fig. 2h and Extended Data Fig. 8a**). The degrees of connection for enhancers were higher than those for promoters (**Extended Data Fig. 8a**), correlating with a high portion of enhancer-enhancer looping (**Fig. 2c**). On average, one enhancer directly interacted with one promoter, and one promoter had direct contacts with almost three enhancers, suggesting a redundant design of robustness for the cis-regulatory network (**Extended Data Fig. 8b**).

To test the functional contribution of direct and indirect enhancer loops to a target promoter in the interaction network, we chose the *CEMIP2* gene locus as a model, which has multiple potential enhancers (annotated as E1-E8) in K562 cells (**Extended Data Fig. 8c**). While E2 interacts directly with the promoter, both E3 and E5 interact strongly with E2, but not with the promoter. Interestingly, deleting any of these three potential enhancers by CRISPR/Case9 resulted in decreased expression of *CEMIP2* (**Extended Data Fig. 8d**). Meanwhile, the *CEMIP2* promoter interactions decreased by deleting any of these elements (**Extended Data Fig. 8e**). These results suggest that deleting one node of the interaction network may affect the stability of the whole regulatory interaction network, resulting in the dysregulation of gene expression.

### HCFC1 and ZNF143 are associated with promoter-centric chromatin looping

We noticed that many loop anchors were not occupied by CTCF or SMC3 (**Fig. 2a**), especially for short-distance loops, suggesting that other factors may organize chromatin looping at these sites. To identify such potential factors, we analyzed the enrichment of 162 transcription factors (TFs) at Hi-TrAC loop anchors in GM12878 and 360 TFs in K562 cells (**Supplemental Table 5**). The top 15 enriched TFs in both cells included CTCF and RAD21 (**Fig. 3a**). Interestingly, the analysis also revealed that both cells shared two highly enriched TFs, HCFC1 and ZNF143, suggesting they may also be broadly involved in orchestrating chromatin looping. HCF1 and ZNF143 are ubiquitously expressed TFs that function at promoters of target genes, regulating cell metabolism, proliferation and differentiation^68–75^. Dysregulation of HCFC1 and ZNF143 is related with the pathogenesis of diseases (e.g. cancer)^76–79^. Accumulating evidence suggest that HCFC1 and ZNF143 may be involved in organizing chromatin structures^8, 51–53, 80–84^. The number of loops co-bound by HCFC1/ZNF143 but not CTCF or RAD21 was similar to the number of loops co-bound by CTCF/RAD21 but not HCFC1 or ZNF143 in GM12878 (**Fig. 3b**). Furthermore, our data indicated that only a small fraction of CTCF/RAD21 co-bound loops were associated with promoters, whereas a striking nearly 90% of promoter loops were anchored by HCFC1/ZNF143 binding (**Fig. 3c-d**). HCFC1/ZNF143 co-bound loops were generally shorter in genomic distance than CTCF/RAD21 loops (**Fig. 3e**). Genes with HCFC1/ZNF143 promoter-promoter loops showed higher expression levels than other genes (**Fig. 3f).** These results suggest a “division of labor” model for chromatin looping by different architectural proteins.

**Fig. 3.**
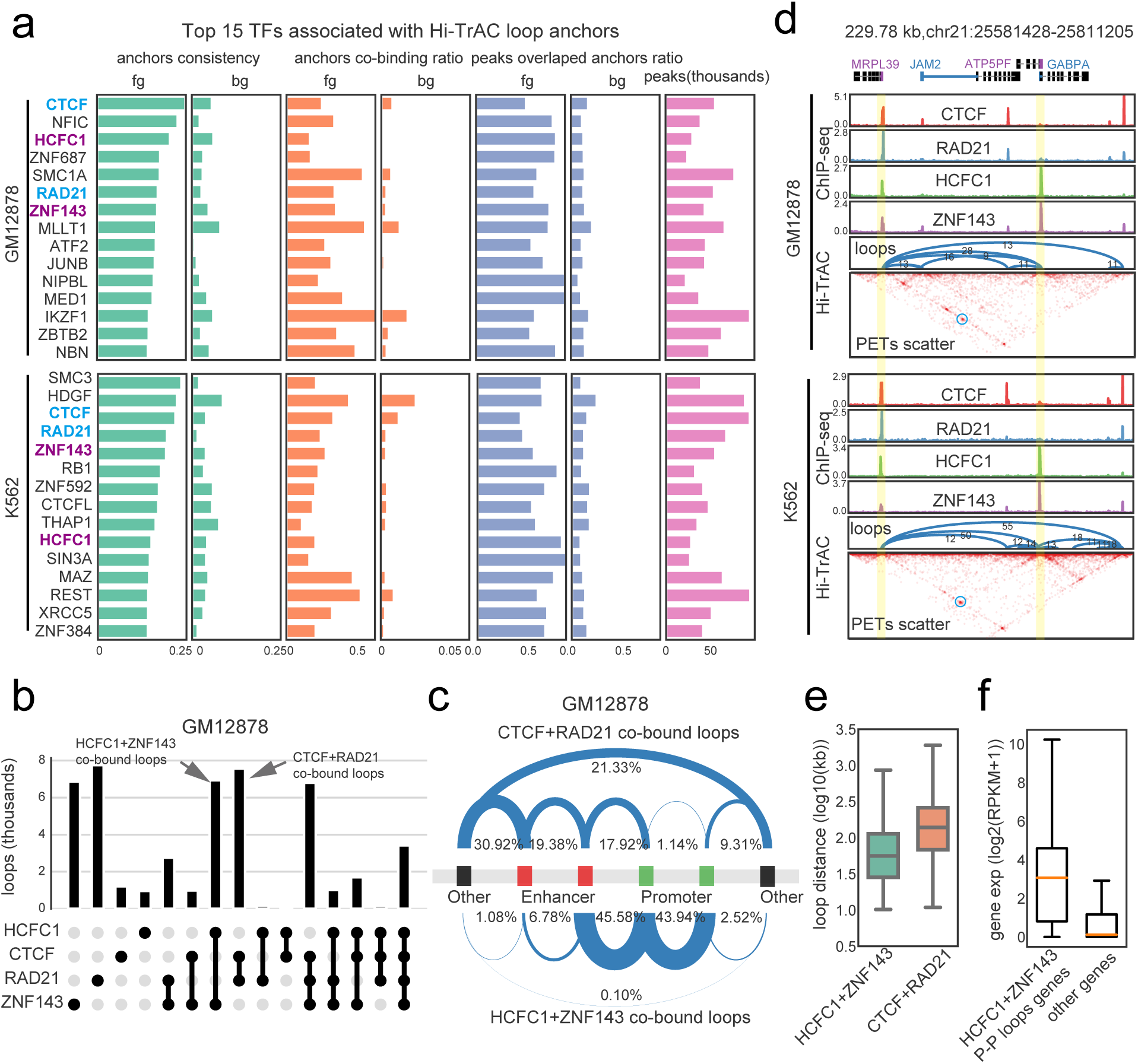
Division of labor in regulating different categories of chromatin loops by distinct transcription factors. **(a)** The top 15 transcription factors associated with the Hi-TrAC loop anchors in GM12878 and K562 cells. The binding sites of 162 and 360 transcription factors were compiled from the ReMap 2020^105^ for GM12878 and K562, respectively. The top 15 factors most significantly associated with loop anchors, sorted by consistency, are shown as indicated on the left of the panel (**Supplemental Table 5).** Overlapped top factors between GM12878 and K562 are highlighted in purple and blue. Fg stands for foreground data, which means the actual loops. Bg stands for background data, which means regions nearby actual loop anchors and used as controls. **(b)** The distribution of loops (top panel) bound by HCFC1, CTCF, RAD21, and ZNF143 alone or in combination (bottom panel) at both anchors in GM12878. **(c)** Summary of loop categories with regard to putative cis-regulatory elements for loops co-bound by HCFC1 plus ZNF143 and loops co-bound by CTCF plus RAD21 in GM12878. **(d)** An example of promoter-promoter loops co-bound by HCFC1 plus ZNF143 but not CTCF plus RAD21. **(e)** The distribution of anchor distance for loops co-bound by HCFC1 plus ZNF143 and loops co-bound by CTCF plus RAD21 in GM12878. **(f)** The expression levels of genes with promoter-promoter loops co-bound by HCFC1 plus ZNF143 and other genes in GM12878.

### Disrupting CTCF, RAD21, ZNF143 and HCFC1 results in distinct perturbation of looping

To test the roles of CTCF, RAD21, HCFC1, and ZNF143 in maintaining chromatin looping, we knocked down (KD) these different factors either individually or in combination in K562 cells (**Extended Data Fig. 9a-b**). The resulting cells were then analyzed with Hi-TrAC (**Supplemental Table 1**). Aggregated enrichment scores of PETs in loop regions compared to nearby regions for all loops were used to compare the effect of knocking down on chromatin looping. While knocking down each of these TFs globally decreased the looping intensity, simultaneously knocking down both CTCF and RAD21 or both HCFC1 and ZNF143 resulted in more severe decreases in chromatin looping (**Supplemental Table 6**, **Fig. 4a**), suggesting that these TFs facilitate looping in general and they may act cooperatively to mediate looping. Consistent with this notion, each knocking down significantly decreased 3,160 (CTCF KD), 3,743 (RAD21 KD), 1,480 (HCFC1 KD) and 1,386 (ZNF143 KD) loops, respectively, while it enhanced smaller numbers of loops: 1,740 (CTCF KD), 934 (RAD21 KD), 779 (HCFC1 KD) and 843 (ZNF143 KD), respectively (**Extended Data Fig. 9c**). Simultaneous knocking down of CTCF and RAD21 decreased 4,249 and enhanced 701 loops, while simultaneous knocking down of HCFC1 and ZNF143 decreased 1,646 and enhanced 734 loops, respectively (**Extended Data Fig. 9c**). The changes of chromatin looping intensity didn’t show a strong correlation with accessibility changes (**Supplemental Table 7, Extended Data Fig. 9d**). We found that 48.8% and 35.5% of the anchors of loops decreased by knocking down CTCF and RAD21, respectively, were other than enhancers and promoters, while lower fractions (24.93% and 25.11%, respectively) of the anchors of loops decreased by knocking down HCFC1 and ZNF143 belonged to this category of accessible regions (**Fig. 4b**). Similarly, the anchors of loops decreased by simultaneous knocking down of CTCF and RAD21 also displayed a higher fraction (36.03%) of non-promoter and non-enhancer regions than that (21.86%) decreased by the simultaneous knocking down of HCFC1 and ZNF143 (**Fig. 4b**). By comparison, knocking down of HCFC1 and ZNF143, either individually or simultaneously, resulted in higher fractions of disrupted enhancer- and promoter-related loops (**Fig. 4b**). The median sizes of loops decreased by knocking down CTCF and/or RAD21 were ∼ 100 kb, while the median sizes of loops decreased by knocking down HCFC1 and/or ZNF143 were ∼20-30 kb (**Fig. 4c**). These results indicate that HCFC1 and ZNF143 are involved in organizing different groups of chromatin loops compared with CTCF with RAD21.

**Fig. 4.**
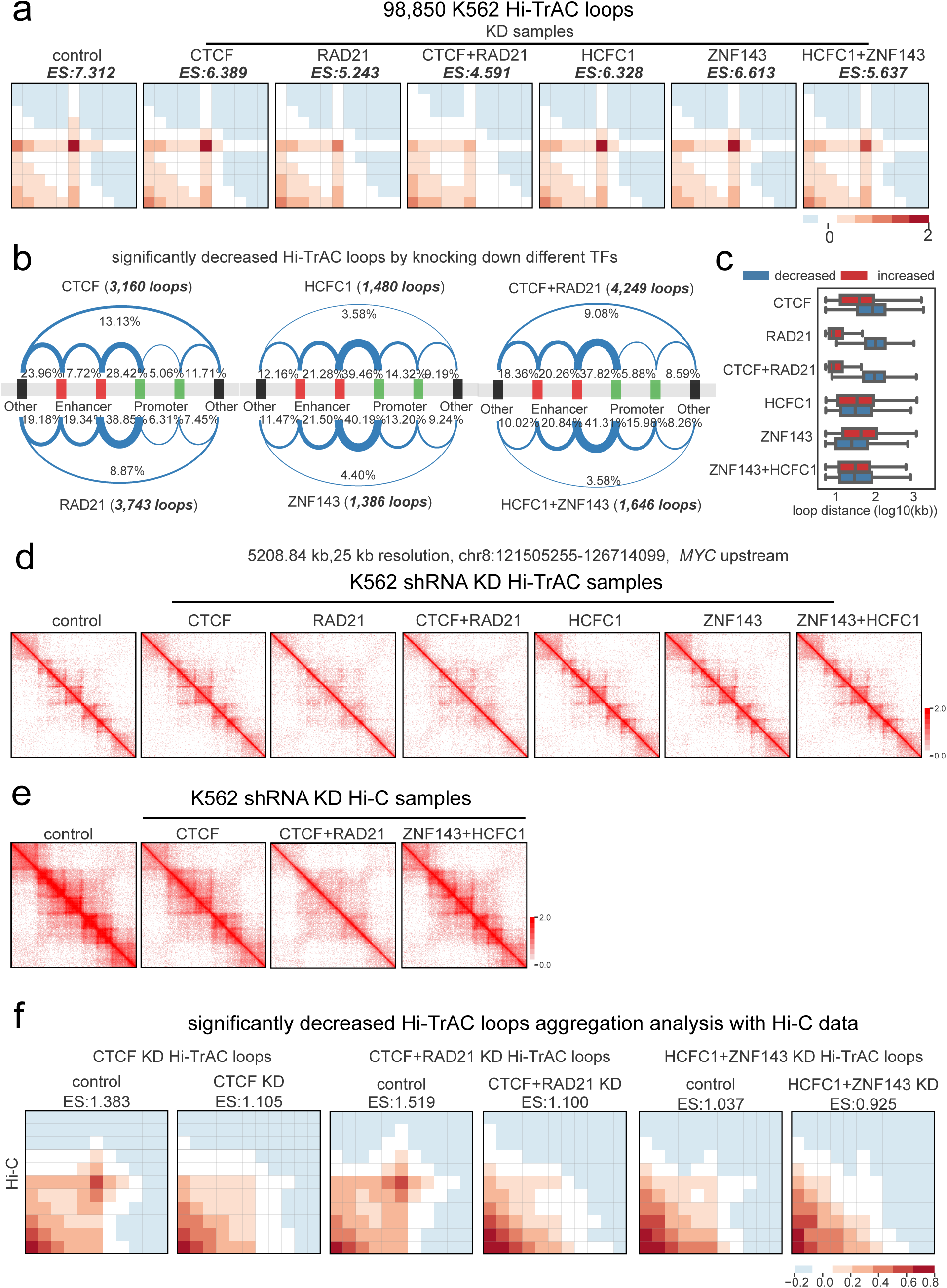
HCFC1 and ZNF143 contribute to organizing chromatin looping. **(a)** Aggregation analysis reveals decreases in chromatin looping intensity after knocking down CTCF, RAD21, HCFC1 and ZNF143 in K562 cells (**Supplemental Table 6**). Enrichment score (ES) is the mean value of all enrichment scores for individual loops. **(b)** Summary of loop categories with regard to putative cis-regulatory elements for decreased loops in the TF knockdown cells. **(c)** The genomic distance distribution of changed loops after TF knockdown. **(d)** An example of domain disruption at the genomic region upstream of *MYC* gene detected by Hi-TrAC after knocking down RAD21 or CTCF with RAD21. **(e)** In situ Hi-C data shows the disruption of chromatin domains in the same region as in panel **d** after knocking down CTCF and RAD21 (**Supplemental Table 8**). **(f)** Loop aggregation analysis for significantly decreased loops detected by Hi-TrAC using the same in situ Hi-C data as in Panel **e**.

To further validate the chromatin structure changes detected by Hi-TrAC, we performed in situ Hi-C with the control and knockdown cells (**Supplemental Table 8**). Consistent with what we observed in Hi-TrAC data (**Fig. 4d**), the in situ Hi-C data also showed severely compromised domain structures by knocking down CTCF and RAD21 but not by knocking down CTCF alone or simultaneously knocking down HCFC1 and ZNF143 (**Fig. 4e**). Hi-TrAC identified significantly decreased loops also showed decreases in the in situ Hi-C data of corresponding cells (**Fig. 4f**). These results demonstrate that Hi-TrAC can accurately detect alterations of chromatin structures.

To investigate the correlation between chromatin looping and gene expression, we analyzed the gene expression profiles in the control and knockdown cells (**Supplemental Table 9**). Knocking down each of these factors resulted in both up-regulated and down-regulated genes: 371 and 239 for CTCF, 926 and 925 for RAD21, 379 and 237 for HCFC1, and 255 and 466 for ZNF143, respectively (**Extended Data Fig. 10a**). Notably, the simultaneous knockdown of HCFC1 and ZNF143 down-regulated many more genes (646) than their individual knockdown. While there were generally no significant correlations between changes in gene expression and chromatin looping, a modestly positive Pearson Correlation Coefficient (PCC) of 0.219 for the down-regulated genes by simultaneously knocking down HCFC1 and ZNF143 was observed (**Extended Data Fig. 10b**), suggesting a possibility that the loops requiring both of these factors contribute to the expression of this group of genes. However, the data also indicated a complex relationship between changes in chromatin looping and gene expression, suggesting that HCFC1 and ZNF143-dependent loops could mediate either gene activation or repression.

### HCFC1 and ZNF143 work separately and synergistically with CTCF and RAD21

To determine if the changes in chromatin looping were direct consequences of depleting these TFs, we checked their chromatin binding profiles by ChIP-seq in knockdown cells (**Supplemental Table 10**). Simultaneously knocking down CTCF and RAD21 drastically reduced the binding of RAD21, whereas only reduced CTCF binding mildly (**Extended Data Fig. 11a**). Interestingly, the binding of HCFC1 was also impaired dramatically. Double knocking down of HCFC1 and ZNF143 significantly impaired their binding, and it also showed mild influence on the binding of CTCF and RAD21 (**Extended Data Fig. 11a**). HCFC1 and ZNF143 binding sites were enriched with the “ACTACANNTCCCA” ZNF143-associated motif (**Extended Data Fig. 11b**). Over 70% of HCFC1 and ZNF143 co-bound sites were located at promoters (**Extended Data Fig. 11c**). In both CTCF with RAD21 and HCFC1 with ZNF143 double knockdown cells, the top enriched motifs in decreased loop anchors included “CTCF” and “GATA” motifs (**Extended Data Fig. 11d**). Even though only 8% of the decreased loop anchors in HCFC1 and ZNF143 double knockdown cells were HCFC1 and ZNF143 co-bound peaks (**Extended Data Fig. 11e**), HCFC1/ZNF143 motif was still one of the top enriched motifs (**Extended Data Fig. 11d**). Furthermore, those decreased loop anchors not bound by HCFC1 or ZNF143 in the HCFC1 and ZNF143 double knockdown cells were enriched with “CTCF” and “GATA” motifs (**Extended Data Fig. 11f**). These binding motif analyses of decreased loop anchors in knockdown cells suggest that HCFC1 and ZNF143 may act both separately and together with CTCF and RAD21.

### HCFC1 and ZNF143 orchestrate gene expression by organizing promoter loops

To further investigate the functions of HCFC1 and ZNF143 in organizing chromatin structures, we examined a primate-specific genomic region which harbored multiple zinc-finger (ZNF) genes. This region exhibited strong promoter-promoter interactions, which is conserved between K562 and GM12878 cells. Multiple HCFC1 and ZNF143 binding peaks were detected at the loop anchors within this locus, whereas no strong CTCF and RAD21 binding was detected (**Fig. 5a**). The simultaneous knockdown of HCFC1 and ZNF143 severely compromised promoter-promoter looping within this region as shown by the Rehoboam plots (**Fig. 5b**), which was accompanied by decreased expression of the target genes (**Fig. 5c**), indicating a critical role of HCFC1 and ZNF143 in regulating looping and the expression of these genes. It was confirmed by ChIP-seq that the decrease in the promoter-promoter looping correlated with impaired bindings of HCFC1 and ZNF143 (**Fig. 5d**). We further validated the disruption of promoter-promoter looping detected by Hi-TrAC in the HCFC1 and ZNF143 knockdown cells using 3C-qPCR assays. The results indicated that both the *ZNF224*-*ZNF284* and *ZNF225*-*ZNF235* loops significantly decreased after knocking down HCFC1 and ZNF143 (**Extended Data Fig. 12a**). Two other randomly selected promoter-promoter loops outside of the ZNF gene cluster region, *MRPL24*-*PRCC* and *NDC1*-*TCEANC2,* were also significantly impaired by knocking down HCFC1 and ZNF143 (**Extended Data Fig. 12a-b**).

**Fig. 5.**
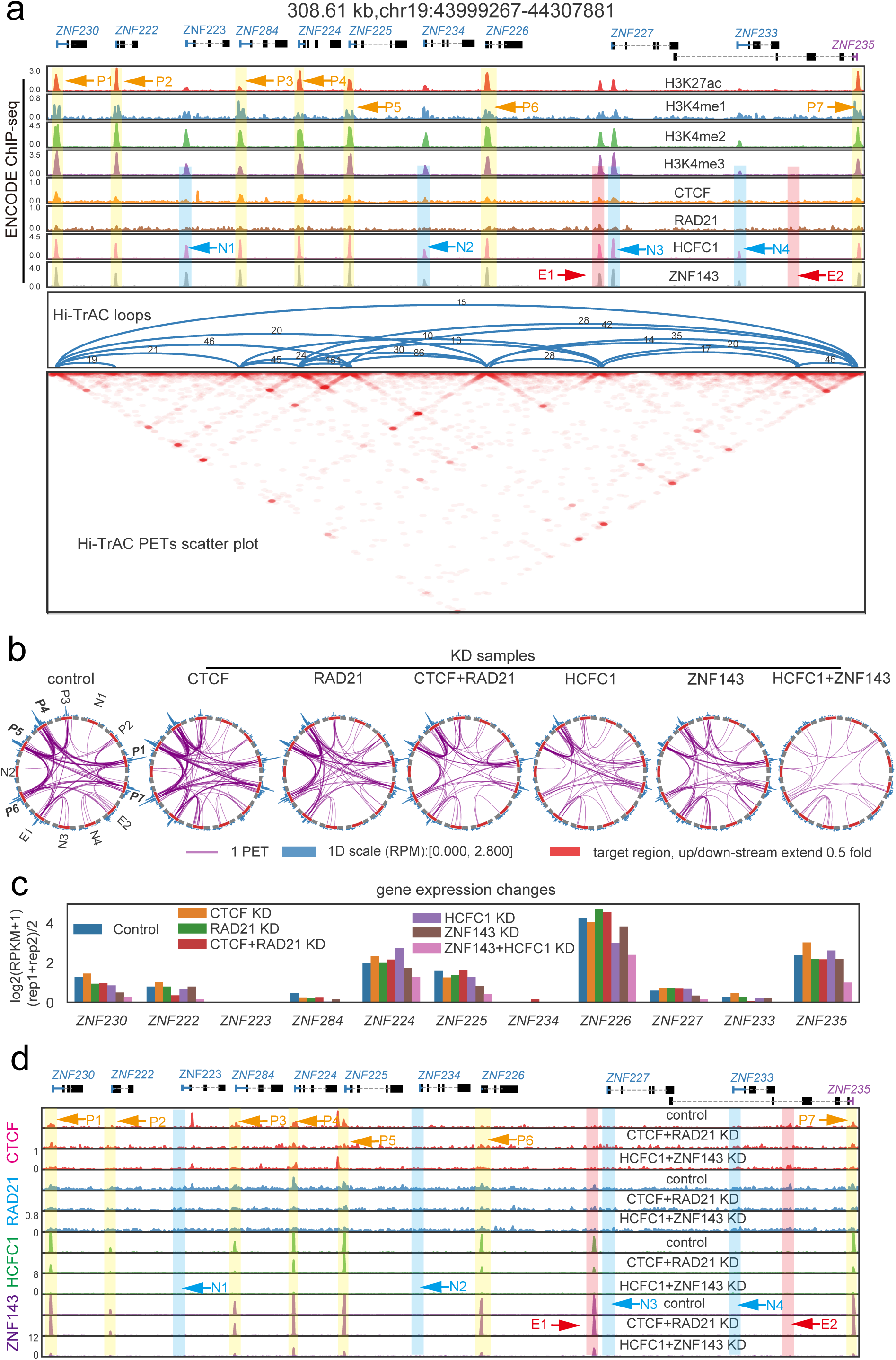
HCFC1 and ZNF143 regulate gene expression through organizing promoter-promoter looping. **(a)** Chromatin loops detected by Hi-TrAC are shown for the ZNF gene cluster on Chromosome 19 in K562 cells. Also shown are ENCODE ChIP-seq signals of active histone modifications and 4 TFs (H3K4me1, 2, 3 and H3K27ac, CTCF, RAD21, HCFC1, and ZNF143). Putative promoters co-bound by HCFC1 and ZNF143 with loops are annotated as P1 to P7, and putative promoters co-bound by HCFC1 and ZNF143 but no loops detected are annotated as N1 to N4. The non-promoter region showing loops and co-bound by HCFC1 and ZNF143 is annotated as E1, and the region showing looping with other region but no HCFC1 and ZNF143 binding is annotated as E2. **(b)** Rehoboam plots of chromatin looping in control and knockdown cells for the ZNF cluster region as annotated in panel **a**. **(c)** The expression changes of the ZNF genes by knocking down CTCF, RAD21, HCFC1, and ZNF143 in K562 cells as measured by RNA-seq. **(d)** Binding profiles of CTCF, RAD21, HCFC1 and ZNF143 at ZNF gene cluster region detected by ChIP-seq in control, CTCF plus RAD21 double knockdown or HCFC1 plus ZNF143 double knockdown K562 cells.

To further test whether the HCFC1 and ZNF143-dependent chromatin loops positively or negatively contribute to the expression of these genes, we deleted either ZNF225 or ZNF234 promoter loop anchor at the ZNF gene cluster region using CRISPR: the expressed *ZNF225* promoter is bound by HCFC1 and ZNF143 and interacts with other promoters in the region, while the non-expressed *ZNF234* promoter has only very weak binding of HCFC1 and ZNF143 and does not interact with other promoters and thus serves as a negative control (**Fig. 5a-c**). Deleting the *ZNF225* promoter loop anchor led to decreased interactions at *ZNF225* promoter and decreased *ZNF225* gene expression as expected (**Extended Data Fig. 12c-d**). Surprisingly, deleting *ZNF225* promoter anchor led to increases in loop formation at other promoters in this region, and also increases in the expression of those genes (**Extended Data Fig. 12c-d**). These results showed that HCFC1 and ZNF143 bound anchors contribute to promoter loop formation, which negatively affects the expression of nearby genes, potentially by directly competing for the limited transcriptional regulators.

## Discussion

We present here Hi-TrAC for mapping the fine architectures of active chromatins. This technique shares the basic concept with Trac-looping, which utilizes Tn5’s ability to integrate DNA bridge into accessible chromatin regions to covalently link physically-contacting chromatin loci, thereby avoiding the proximity ligation that is required for the 3C-derived techniques^59^. Trac-looping needs 50-100 million cells and 5-7 days of bench work. Hi-TrAC is much more versatile and efficient, can be performed with as few as 0.01 million cells and less than two days of bench work. Moreover, it captures detailed interaction information with a resolution up to 200 bp with only 300 million sequencing reads, detecting interactions between regulatory elements with high sensitivity. Chromatin interactions detected by Hi-TrAC are highly correlated with those detected by Micro-C, MCC and even CRISPRi, suggesting that Hi-TrAC can reliably capture spatial contacts of regulatory elements. Moreover, comparing to Micro-C, Hi-TrAC detected four times more chromatin loops (300k vs 64k) with only 10% of PETs (223M vs 2,396M); Hi-TrAC detected almost all the interactions between potential enhancers and their target promoters as detected by MCC. Comparing to other 3C-based techniques, Hi-TrAC also showed advantage in detecting much more chromatin loops at high resolution with less sequencing amount. Thus, Hi-TrAC can serve as an inexpensive, highly sensitive and robust alternative method for analyzing interactions among regulatory elements of chromatin and transcription.

The bridging linker used in Hi-TrAC works like a ruler charting the genome map. To be bridged by the linker, the spatial distance between two interacting loci should be shorter than the length of the linker. We tested linkers with different lengths and found that a linker with a 30 bp spacer between the flanking Tn5 binding sites performed best. Longer linkers captured too many inter-chromatin interactions, whereas shorter linkers lost many distal interactions (**Supplemental Table 11**). The chosen bridging linker is estimated as ∼20 nm long, suggesting the spatial physical distance between Hi-TrAC captured interacting regions are within that range.

Enhancer-promoter interaction loops play a critical role in controlling the temporospatial expression of genes^31, 33, 35, 36, 38, 40, 43^. Taking the β-globin locus as an example, the expression switching of fetal and adult hemoglobin is regulated by interactions between a locus control region (LCR) and promoters of corresponding genes with the help of transcription factors^85, 86^. To gain comprehensive information on the mechanisms of gene regulation genome-widely, it is important to identify the regulatory interaction networks of enhancers and promoters. However, even the most comprehensive 3D data in previous studies only provide limited information on genome-wide enhancer-promoter loops^8, 61^. To our knowledge, we now have provided the most comprehensive interaction data among accessible chromatin regions in GM12878, K562 cells and mESCs, reporting a total of about 500,000 chromatin loops. Further, our data also revealed shared chromatin loops and cell-specific loops that contribute to cell-specific expression of genes that are responsible for differentiation and cellular function. Even for genes showing similar expression levels in both cell types, the interactions of regulatory elements, especially of enhancers, exhibit different patterns, suggesting that gene expression could be differentially regulated in different cells.

Chromatin loops can be roughly separated into three categories based on the location of loop anchors relative to chromatin domain organization: (1) in the TAD boundaries; (2) in the sub-TAD boundaries; and (3) within TADs and sub-TADs. They can also be categorized based on the functional annotation of the loop anchors: (1) promoters; (2) enhancers; and (3) others. While the loops linking enhancers and promoters are often found within TADs or sub-TADs, others are mapped to the boundaries of these chromatin domains. Generally, the loops originating from the domain boundaries are much longer (>100 kb) than those linking enhancers and promoters (<100kb). Previous data have established that CTCF and cohesin complex play critical roles in maintaining the chromatin interaction of TADs^48, 50, 87–89^ via an extrusion model^18–23^. Several specific transcription factors have been found to contribute to chromatin looping between enhancers and promoter^51–58^. YY1 facilitates the loop formation between enhancer and promoter and regulates gene expression, and ZNF143 was reported to act with CTCF to facilitate chromatin looping. However, in general, the mechanisms that regulate enhancer-promoter looping requires much clarification. In this study, we found that in addition to CTCF and RAD21, ZNF143 and HCFC1 are also among the top enriched factors and shared among GM12878 and K562 cells, suggesting that these two factors may play a general role in chromatin looping. Interestingly, over 96% of the loop anchors co-bound by ZNF143 and HCFC1 in GM12878 cells involved enhancers and promoters, while only 38% of the loop anchors were co-bound by CTCF and RAD21 in GM12878 cells involved enhancers and promoters, suggesting that ZNF143 and HCFC1 play a general role in contributing to enhancer-promoter loops. The loops disrupted by knocking down HCFC1 and/or ZNF143 showed a median size of 20 kb, consistent with the size of enhancer and promoter interactions, while the loops disrupted by knocking down CTCF and/or RAD21 showed a median size of about 100 kb, consistent with the distance between domain boundaries.

Although it was previously suggested that promoter-promoter interactions may facilitate gene expression by bringing the target promoters in close proximity^29, 90–92^, direct data to support the hypothesis are needed. Here, we found that HCFC1 and ZNF143 bound to the anchors of promoter loops at the ZNF gene cluster locus; the simultaneous knocking down HCFC1 and ZNF143 disrupted promoter-promoter loops and decreased the expression of these target promoters and thus providing strong evidence that the promoter-promoter looping contributes to the gene expression. Based on these results, we propose that different architectural proteins show a division of labor for organizing chromatin looping: CTCF and RAD21 are responsible for building the outer frame of chromatin domains, whereas HCFC1 and ZNF143 decorate the inner structures by organizing looping between regulatory elements. With the development and completion of high-resolution genome structure and more chromatin binding datasets of transcription factors, more chromatin looping regulators will be identified, including general ones and cell type-specific ones. Our comprehensive data on the interaction network among accessible chromatin regions provide a rich resource to further explore the complex function and mechanisms in genome organization.

## Methods

### Hi-TrAC experimental procedures

Cells were fixed with 1% Formaldehyde in culture medium at room temperature for 10 minutes. Wash the cells twice with 1 mL ice-cold PBS, then keep cells on ice. The Tnp complex was assembled by mixing 2 μL short adapter (50 μM), 2 μL bridging linker (25 μM) (Supplemental Table 12), 2 μL glycerol and 4 μL Tn5 (100 μM), then incubated at room temperature for 20 minutes. Resuspend cells with 100 μL reaction buffer (50 mM Tris-acetate, pH 7.5, 150 mM potassium acetate, 10mM magnesium acetate, 4 mM spermidine, 0.5% NP-40), and incubate at room temperature for 10 minutes. Add 10 μL Tnp complex to the permeabilized cells, then mix gently by pipetting and incubate on a 37 °C thermomixer for 4 hours with interval mixing. The reaction was stopped by adding EDTA (25 mM final concentration) and SDS (0.3% final concentration). Add 2 μL protease K (20 mg/mL) to the reaction mixture and incubate at 65 °C overnight to reverse crosslinking. DNA was purified by Phenol-Chloroform extraction. The gaps in DNA were repaired by T4 DNA polymerase in the reaction mixture containing dNTPs at room temperature for 30 minutes. The free bridging linker (68 bp) was removed by selectively binding large DNA fragments (>100 bp) to AMPure XP beads. The DNA was eluted from AMPure XP beads with 80 μL elution buffer and digested with the restriction enzymes 2 μL MluCI (NEB, R0538L) and 2 μL NlaIII (NEB, R0125L) in 100 μL reaction mixture at 37 °C for 30 minutes. The reaction mixture was adjusted to 1x B&W buffer by adding 100 μL 2x B&W buffer (10 mM Tris-HCl, pH 7.5, 1 mM EDTA, 2 M NaCl, 0.1% Tween-20) and then mixed with 5 μL Streptavidin C1 beads (Invitrogen, 65001) for 30 minutes with rotation. The beads were washed for 5 times with 1 mL 1x B&W buffer. Biotin-labeled DNA fragments captured on beads were ligated to multiplexing adaptors by adding 5 μL each adapter (50 μM) and 1 μL T7 DNA ligase (NEB, M0318L) in 100 μL ligation mixture and incubating at room temperature for 1 hour with rotation. Before PCR amplification, the beads were washed 5 times with 1x B&W buffer. The Hi-TrAC libraries were then amplified with multiplexing indexed primers in the following reaction mixture: 20 μL Phusion HF PCR Master Mix (NEB, M0531S), 1 μL Illumina Multiplexing PCR primer 1.0 (10 μM), 1 μL Illumina Multiplexing PCR index primer (10 μM) and 18 μL H_2_O for 12 cycles. The DNA fragments between 300 bp and 700 bp were excised for paired-end sequencing on Illumina platforms.

### Knockdown and western blotting

Knockdown of CTCF, RAD21, HCFC1 and ZNF143 in K562 cells was performed by transduction with shRNA lentivirus. shRNA templates were cloned into pGreenPuro lentivector (System Biosciences, SI505A-1). shRNA targets are: Control-shRNA-1: GCGCGATAGCGCTAATAATTT, Control-shRNA-2: CAACAAGATGAAGAGCACCAA; CTCF-shRNA-1: GGAGAAACGAAGAAGAGTA, CTCF-shRNA-2: GTAGAAGTCAGCAAATTAA; RAD21-shRNA-1: AGAGTTGGATAGCAAGACA, RAD21-shRNA-2: GGAAGCTAATTGTTGACAGTGTCAA; HCFC1-shRNA-1: GCAACCACCATCGGAAATAAA, HCFC1-shRNA-2: AGAACAACATTCCAAGGTACCTGAA; ZNF143-shRNA-1: GCTACAAGAGTAACTGCTAAA, ZNF143-shRNA-2: GGACGACGTTGTTTCTACACAAGTA. Co-transfect 12 μg lentivector with packaging plasmids 9 μg psPAX2 (Addgene, #12260) and 3 μg pMD2.G (Addgene, #12259) into 293T cells cultured in 100 mm dish. Change with 12 mL fresh medium at 12 hours after transfection, and medium supernatant containing virus was collected at 72 hours after transfection. Add the medium to 3 million K562 cells to start transduction. Cells were harvested 72 hours after infection. Protein expression was checked by western blotting. We used Invitrogen NuPAGE gel electrophoresis system following owner’s manual, and proteins were transferred onto PVDF membrane. Primary antibodies used for detecting corresponding proteins are: anti-CTCF (Cell Signaling Technology, 3418S), anti-RAD21 (Abcam, ab217678), anti-HCFC1 (Santa Cruz Biotechnology, sc-390950) and anti-ZNF143 (Abnova Corporation, H00007702-M01).

### RNA-seq library construction

Total RNA from 5,000 cells were extracted and purified with QIAzol Lysis Reagent (QIAGEN) and RNeasy mini kit (QIAGEN). RNA-seq library was constructed with purified RNA following the Smart-seq2 protocol^93^.

### Generating loop anchor deletion cells

CRISPR targeting sequences were designed and cloned into pSpCas9(BB)-2A-Puro vector (Addgene #62988). Following transfection of K562 cells with Cas9 and sgRNA expressing plasmids for 24 hours, the cells were treated with 2 μg/mL Puromycin for 48 hours to kill non-transfected cells. Surviving cells were sorted into 96-well plate at a density of one cell per well, cultured for two to three weeks and genotyped using specific PCR primers for identification of loop anchor deletion clones. The targeting sequences used are following: CEMIP2-E2, 1-GATCGAGTTCTAGTTGACCC, 2-GTGCGTCTATGAATCTGCGC; CEMIP2-E3, 1-GTAAGCACATGGCCCGTCAG, 2-TCGAACAGGAACGTACTATC; CEMIP2-E5, 1-CTAACGCAATCCACCTAGAA, 2-TAAGGCTCTCTACTTAGCGG; ZNF225-promoter, 1-TGGCGCTTAACGACGAACCC, 2-TTTATGGGGCACGGCGACCA; ZNF234-promoter, 1-AAGGAGGATCCTATACGTGA, 2-TAAGCCGCAACGTGACTCTG.

### Public data and pre-processing

Public data used in this study, including Hi-C, HiChIP, ChIA-PET, capture Hi-C, MCC, Micro-C, RNA-seq, ATAC-seq, DNase-seq, and ChIP-seq, were summarized in **Supplemental Information**. Biological and technical replicates were merged for the same factor and only unique reads were used for the following analyses.

### Public genomic annotations

Human (gencode.v30.basic.annotation.gtf) and mouse (gencode.vM21.basic.annotation.gtf) gene annotations from GENCODE^94^ were used in any gene-related analysis. Human genome version hg38 and mouse genome version mm10 were used in this study. If human or mouse data are generated in other genome versions, they are always converted to hg38 or mm10 for analysis.

Putative enhancer and promoter annotations of human cells were obtained from NIH Roadmap Epigenomics Consortium^96^ and processed as following: 1) all regions annotated as “Enh” and “Tss” were collected; 2) overlapping regions were merged with BEDtools merge; 3) merged regions were further annotated by annotatePeaks.pl in HOMER package^97^ with gene annotation file downloaded from GENCODE^94^; if a region is within 2kb either upstream or downstream of a TSS, it is defined as a promoter; otherwise it is defined as an enhancer. 4) neighboring enhancers or promoters with gaps < 100 bp were merged again by BEDtools merge.

Putative cis-regulatory elements of mouse embryonic stem cells were defined by ATAC-seq peaks. If a peak is within 2 kb either upstream or downstream of a TSS, it is defined as a promoter; otherwise, it is defined as an enhancer.

### Pre-processing of Hi-TrAC data

Raw paired-end reads in FASTQ files were first trimmed of the linker sequence CTGTCTCTTATACACATCT from both ends. Only paired-end tags (PETs) with both ends with a length ≥ 10bp were kept. Trimmed PETs were mapped to hg38 using Bowtie2^98^ with --end-to-end --very-sensitive parameter. Mapped PETs with MAPQ ≥ 10 were converted to BEDPE files. Mapped PETs with a distance shorter than 1 kb without linker sequence in any end were further filtered. PCR replicates of PETs were filtered if the locations of both ends were identical. Unique intra-chromosomal PETs (cis PETs) as BEDPE files were mainly used for downstream analysis. All the described processing steps were summarized as tracPre2.py in the cLoops2 package^99^. Quality control statistical results were also generated by tracPre2.py. BEDPE files were used to analyze PET level properties, and the cLoops2 pre module processed them to cLoops2 data directories for other analyses such as domain-calling, loop-calling, and visualization.

### Virtual 4C signals of Hi-TrAC or Micro-C

Virtual 4C signals were generated by only keeping the PETs with one end located within 1kb either upstream or downstream of target TSSs. These PETs were then piled up as the 1D signal. The method is implemented in the cLoops2 plot module for visualization or the cLoops2 dump module for data extraction^99^.

### Comparisons between mESC Hi-TrAC and Micro-C loops

Micro-C loops were called by HiCCUPS in Juicer package (v1.22.01)^100^ with parameter settings of –cpu –ignore-sparsity-r 2500 -f 0.1 -k KR -p 4 -i 8 -d 2 for the 2.6 billion PETs HIC file downloaded from GEO, according to the original paper as leading to the most of loops compared to other resolutions. More loops were called from the Micro-C data than the original paper due to the upgrades of the HiCCUPS algorithm^60^. The loop calling algorithm described in cLoops^101^ was slightly improved and implemented as the cLoops2 callLoops module^99^ for loop-calling with Hi-TrAC data. mESC Hi-TrAC loops were called by the cLoops2 callLoops module with parameters of -eps 200,500,1000,2000 -minPts 20 -p 30 -w -j -i -max_cut -cut 5000. For the overlapping analysis, loop anchors were extended to 5 kb, unique and overlapped loops were obtained by pairtopair subcommand in BEDTools (v2.29.2) package^102^ with -type notboth or - type both options.

### Calling loops from Hi-TrAC data of human cells

The cLoops2 callLoops module with key parameters settings of -eps 200,500,1000,2000 -minPts 10 -max_cut was used to call loops from GM12878 Hi-TrAC data, requiring a loop supported by at least 10 PETs. For K562 Hi-TrAC data, loops were called by parameters settings of -eps 200,500,1000,2000 -minPts 10 -cut 5000 to filter PETs with distance short then 5kb, by which the default parameters will automatically filter all loops short than 20 kb.

### Loop aggregation analysis

An 11 x 11 contact matrix was constructed for a loop from interacting PETs, together with its five upstream and downstream windows of the same size. An individual enrichment score for a loop was calculated as the number of PETs in the 11 x 11 contact matrix center divided by the mean value of all others. The global enrichment score was the mean value of all enrichment scores for individual loops. The 11x 11 contact matrix was further normalized by the total number of PETs in the matrix and z-score normalization. Heatmap was plotted of the average matrix for all normalized 11 x 11 matrices. The analysis was implemented in cLoops2 agg module with the option of -loops. Except for the parameters specifically mentioned, the default parameter with -loop_norm was used to generate visualization results.

### CTCF motif orientations

The whole-genome-wide CTCT motif orientations were annotated by FIMO^103^ with CTCF motif recorded in CIS-BP database^104^.

### Calling differential loops

Loops from samples under different conditions were combined and quantified in both conditions. The neighboring regions nearby loop anchors, which were the permutated nearby background regions defined in cLoops for estimation loops statistical test, were also quantified. The background data for the two conditions were fitted linearly. The fitted linear model was used to transform the PETs in loops of treatment dataset to the control set, assuming there should be no difference in background data. False discovery rate (FDR) is a required parameter to find the cutoffs of average and fold change in the background data MA plot. The cutoffs were then applied to the transformed loops data. Poisson p-values were finally assigned to each loop as following,

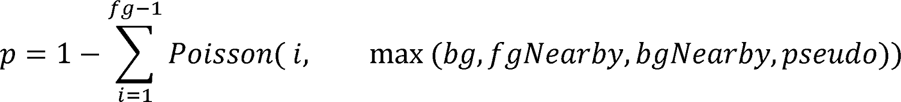

Where fg stands for the bigger value of PETs in the testing loop for treatment vs. control, bg stands for the smaller value for comparison, fgNearby is the number of PETs for background data for the testing condition, and bgNearby is the number of PETs for background data for the control condition. Pseudo is a general noise control value, set to 1 for all times. All numbers except pseudo here are transformed by the linear fitting above the background data. The P-values were corrected by Bonferroni correction, and by default 0.01 was used as the cutoff for significance. This algorithm was implemented in the cLoops2 callDiffLoops module^99^, and differentially enriched loops were called with key parameters of -fdr 0.05 for GM12878 vs. K562 Hi-TrAC data.

### Transcription factors associated with Hi-TrAC loops

Integration of public ChIP-seq data and Hi-TrAC loops was intended to identify transcription factors associated with chromatin looping. We collected the binding sites for 162 transcription factors in GM12878 and 360 transcription factors in K562 from ReMap 2020 ^105^ (remap2020_all_macs2_hg38_v1_0.bed) on 2020-08-09. For a loop, both the regions of three sizes upstream of its left anchor (smaller coordinate in the genome) and downstream of its right anchor (bigger coordinate in the genome) were linked as background data (false loops) for comparisons, to calculate the enrichment of TF binding sites on both anchors. Any background regions overlapping with true loop anchors were removed. The overlaps between anchors and background regions with transcription factor binding sites were compiled into the left anchors’ matrix and the right anchors’ matrix. Rows are anchors or backgrounds, and columns are factors in the binary matrix. In the binary matrix, 1 indicates the anchor is bound by the factor and 0 stands for no binding. With the two binary matrices, the following attributes were calculated and used to find enriched transcription factors: 1) anchors consistency: for a TF, a vector from the left anchor matrix and a vector from the right anchor matrix were used to calculate the Spearman correlation coefficient, indicating the co-binding consistency of a factor at both anchors. 2) anchors co-binding ratio: for a TF, the ratio of anchors bound by the TF. 3) TF peaks overlap ratio: for a TF, ratio of peaks overlapping with loop anchor regions. To filter TFs, the following cutoffs were used: 1) comparing to the background, ratio of consistency > 2; 2) anchors co-binding ratio >0.1; 3) comparing to the background, the ratio of co-binding ratio >2; 4) comparing to the background, the ratio of TF peaks overlap ratio ≥ 1. Except for using the anchor flanking regions as background, we also implemented the random shuffling value 1000 times as background to ensure the observed attributes are higher than permutation background and require FDR < 0.001. The remaining TFs were sorted by consistency in descending order, by which known looping associated factors such as CTCF and cohesin are among the top-ranked factors.

### Analysis of the Hi-TrAC data from the control and TF knockdown K562 cells

Raw Hi-TrAC data were processed by tracPre2.py first to extract the unique cis PETs (**Supplemental Table 1**). Aggregated loops analysis was performed to obtain the enrichment scores of all loops called from K562 Hi-TrAC data and to check the consistency among two shRNAs and biological replicates. All unique cis PETs from the same TF knockdowns were pooled together and sub-sampled to 74 million (the pooled control sample has the least PETs of 74.37 million) for all downstream analyses. The global enrichment scores were also used to show the global changes in all or subsets of K562 Hi-TrAC loops in the TF knockdown samples. Differentially enriched loops from the knockdown samples to control samples (all loops called from K562 Hi-TrAC data were used as the comparing set) were called with key parameters of - noPCorr -pcut 0.001, by which the above-uncorrected Poisson test *P*-value 0.001 was used to select significantly changed loops.

### Rehoboam plots for visualization of interaction changes

In a Rehoboam plot, each putative cis-regulatory element inferred from Hi-TrAC loops or other sources with extended nearby regions was shown as a part of a circle, Hi-TrAC 1D profiles were shown outside the circle, and Hi-TrAC interaction densities were shown as the widths of arches or each Hi-TrAC PET shown as an arch among circle parts. Viewpoints can be set only to keep the PETs oriented from some of the elements. We name the visualization result as a Rehoboam plot because it looks like a predicted divergence from the AI system named Rehoboam in WESTWORLD Season Three. This visualization method is implemented in the cLoops2 montage module^99^.

### Analysis of Hi-C data from the control and TF knockdown K562 cells

Raw Hi-C data reads were processed to human reference genome hg38 by HiC-Pro (v2.11.1)^106^. Only intra-chromosomal PETs from the files with a suffix of allValidPairs output by HiC-Pro were used for all following analyses. The cLoops2 plot module was used to generate visualization plots. Replicates of the Hi-C libraries were combined to validate the decreased loops detected by Hi-TrAC with aggregation analysis, and only the loop and nearby region with more than 20 PETs were used to do the analysis considering the sparsity of Hi-C interacting PETs.

### Analysis of RNA-seq data from the control and TF knockdown K562 cells

Raw RNA-seq data reads were mapped to human reference genome hg38 by STAR (v2.7.3a)^107^. Gene annotation file (v30) downloaded from GENCODE ^94^ was used to quantify gene expression level by Cufflinks (v2.2.1)^108^. Further, significant differentially expressed genes (knocking down samples vs. control) were called by Cuffdiff (v2.2.1) in Cufflinks package, requiring *P*-value < 0.001 and fold changes ≥1.

### ChIP-seq

Control and TF knockdown K562 cells were fixed with 1% Formaldehyde in culture medium at room temperature for 10 minutes. Wash the cells twice with 1 mL ice-cold PBS, then keep cells on ice. 0.1 million fixed cells were used for CTCF and RAD21 ChIP-seq library preparation, and 0.5 million cells were used for HCFC1 and ZNF143 ChIP-seq library preparation. Resuspend cells with 1x TE buffer provided with 1 mM PMSF and 1x protease inhibitor cocktail. Chromatin shearing was performed on a Diagenode Bioruptor Pico sonication device at 4 °C for 6 cycles with 30 sec on and 30 sec off, resulting in 200-1000 bp fragments. Adjust chromatin solution to 1x RIPA buffer ( 1x TE, 0.1% SDS, 0.1% Sodium Deoxycholate and 1% Triton X-100) plus 200 mM NaCl. Collect the chromatin supernatant after centrifugation at 13,000 rpm for 10 min in a 4°C microcentrifuge. 10% of the chromatin supernatant was saved as input. Mix 2 μg antibody with 20 μL Dynabeads Protein A beads (ThermoFisher Scientific, Cat. 10001D) with rotation at room temperature for 1 hour. Wash the beads once with 1x PBS, then add chromatin solution to beads, and incubate at 4 °C overnight with rotation. Wash the beads twice with RIPA buffer, then twice with RIPA buffer plus 300 mM NaCl, then twice with LiCl buffer (1x TE, 250 mM LiCl, 0.5% NP-40, 0.5% Sodium Deoxycholate), and finally twice with 1x TE buffer. Elute DNA and reverse crosslinking by protease K digestion and incubating at 65 °C for 6 hours. Purify DNA with MinElute Reaction Cleanup Kit (QIAGEN, Cat. 28206). Repair ends of DNA using End-It DNA End-Repair Kit (Lucigen, Cat. ER0720), then perform A-tailing with Klenow Fragment (3’-> 5’ exo-) (NEB, Cat. M0212S) provided with dATP, and then perform adapter ligation with T4 DNA ligase (NEB, Cat. M0202L). Amplify the library and add index by PCR. DNA fragments between 200 bp and 600 bp were purified and sequenced on Illumina platforms.

The following antibodies were used in the experiments: anti-CTCF (Cell Signaling Technology, Cat. 3418S), anti-RAD21 (Abcam, Cat. ab217678), anti-HCFC1 (Cell Signaling Technology, Cat. 69690S) anti-ZNF143 (Abnova, Cat. H00007702-M01).

### ATAC-seq

ATAC-seq was performed with 50,000 cells following the protocol as reported^109^.

### 3C-qPCR

Resuspend 1 million fixed cells with 1 mL ice-cold lysis buffer ( 10 mM Tris-HCl, pH 8.0, 10 mM NaCl, 0.2% NP-40, 1 × protease inhibitor), then incubate on ice for 20 minutes. Collect cells by centrifugation at 4 °C, then resuspend with diluted CutSmart buffer ( 346 μL H_2_O, 50 μL 10 × CutSmart Buffer, 44 μL 1% SDS). Incubate at 65 °C for 10 minutes. Add 50 μL of 10% Triton X-100, then shake on a thermomixer at 37 °C for 1 hour. Add 100 U selected restriction enzyme (for *ZNF224*-*ZNF284*, use BamHI; for *ZNF225*-*ZNF235*, use HindIII; for *MRPL24*-*PRCC* and *NDC1*-*TCEANC2*, use NcoI), then incubate at 37 °C overnight with shaking. Collect digested nuclei by centrifugation, then resuspend with 100 μL inactivation buffer (1 × PBS, 1% SDS), and incubate at 65 °C for 20 minutes. Add 895 μL diluted T4 ligation buffer (695 μL H_2_O, 100 μL 10 × T4 DNA Ligase Reaction buffer, 100 μL 10% Triton X-100) and mix well. Add 100 U T4 DNA ligase, and incubate at 16 °C overnight. Add 30 μL 10% SDS and 100 μg Protease K to stop the ligation reaction, then incubate at 65 °C to reverse crosslinking. Purify 3C libraries by Phenol-Chloroform extraction. Design primers and probes according to the sequence of the interaction pair, then quantify the interaction frequency by qPCR.

### Analysis of ChIP-seq and ATAC-seq data from the control and TF knockdown K562 cells

Raw ChIP-seq and ATAC-seq data reads were mapped to human reference genome hg38 by Bowtie2^98^ with key parameters of --local --very-sensitive --no-unal --no-mixed --no-discordant. Mapped PETs with MAPQ >=10 were converted to normalized signals (reads per million) as bigWig files by deepTools^110^ for visualization or aggregation analysis.

### Motif analysis

Motif analysis for Hi-TrAC loop anchors or ChIP-seq peaks was performed by findMotifsGenome.pl in HOMER package^97^. Only top-ranked significant known motifs were shown.

### GO terms enrichment analysis

GO terms enrichment analysis for genes was performed by script findGO.pl in HOMER package^97^, requiring more than ten overlapping genes in the terms, and there are fewer than 1000 genes in the terms. Only top enriched terms sorted by *P*-values were shown.

### Data visualization

Most of 1D profile and heatmap visualizations were shown by the cLoops2 plot module. Networks were visualized and analyzed by NetworkX^111^. Other plots were generated by matplotlib^112^ and seaborn^113^.

### Code availability

Major analyses are summarized in the cLoops2 package and freely available at GitHub with documentation, test data and updates: https://github.com/YaqiangCao/cLoops2.

### Data availability

Hi-TrAC, RNA-seq, ATAC-seq, Hi-C, and ChIP-seq data generated by this study have been deposited to GEO with accession of GSE180175.

## ACKNOWLEDGEMENTS

We thank the NHLBI DNA Sequencing Core Facility for sequencing the libraries; and the NHLBI Flow Cytometry Core facility for sorting the cells. We thank Dr. Warren Leonard for critical reading of the manuscript. The work was supported by Division of Intramural Research, National Heart, Lung and Blood Institute and the 4DN Transformative Collaborative Project Award (A-0066) (KZ).

## AUTHOR CONTRIBUTIONS

K.Z. conceived the project. S.L. designed and performed the experiments. Y.C. analyzed the data. K.C. contributed to the experiments and Q.T. contributed to the experimental design. S.L., Y.C., and K.Z. wrote the paper.

---

**Extended Data Fig. 1.**
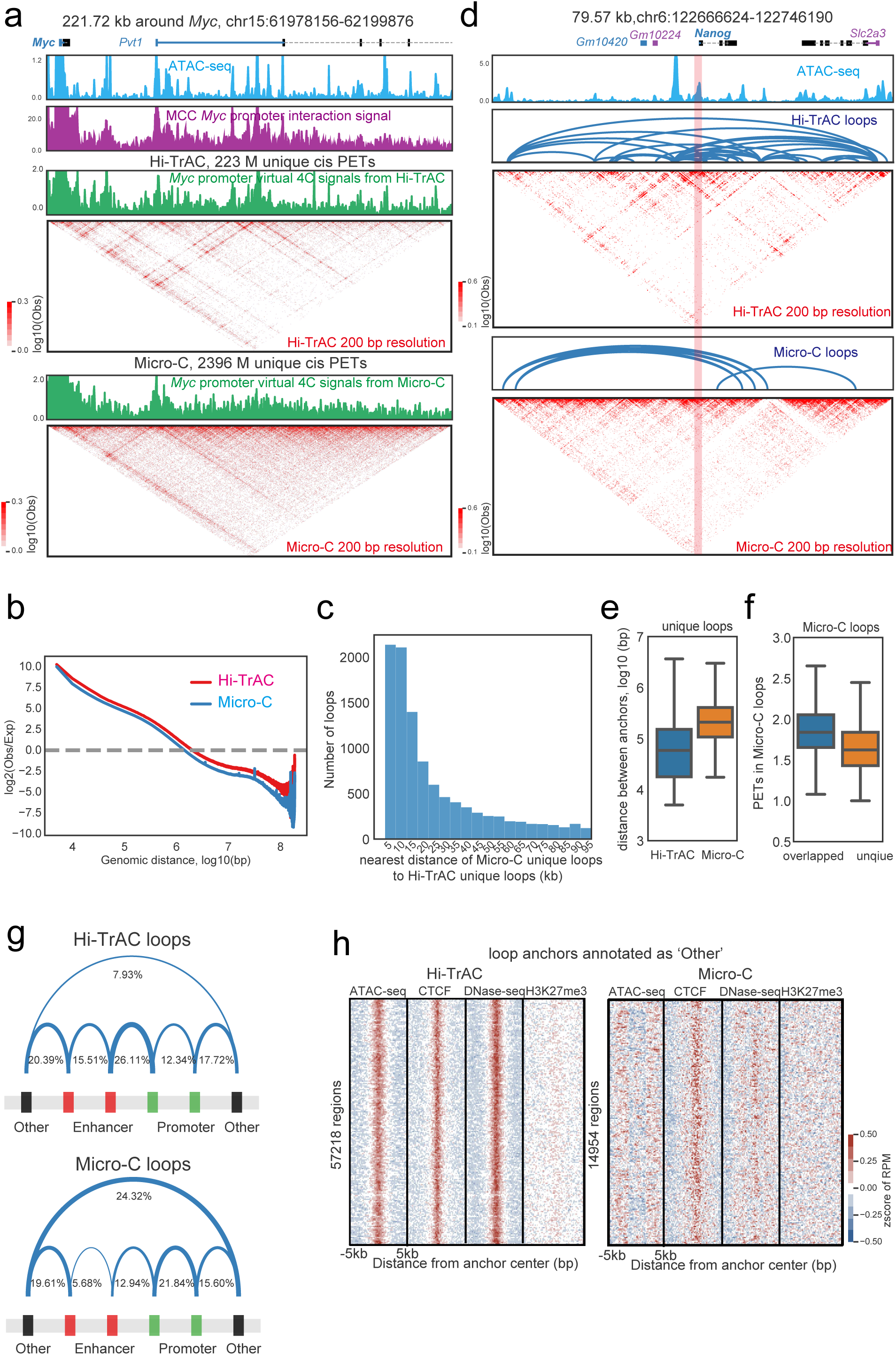
Comparison of the performance in detecting chromatin loops between Hi-TrAC and Micro-C. **(a)** Comparison of interactions in *Myc* gene and its immediate downstream region. The interaction heatmaps from the Hi-TrAC and Micro-C data were shown at 200 bp resolution. **(b)** Distribution of interacting PETs frequency against genomic distance for Hi-TrAC and Micro-C. Both axes are log10 transformed. The random shuffling of two ends of the PETs 10 times was used to generate expected background. The analysis was performed with the cLoops2 estDis module. **(c)** Distance distribution of unique loops from 2.6 billion Micro-C data to unique loops of Hi-TrAC. More than 4,000 (extra ∼6%) Micro-C loops are not overlapped with Hi-TrAC loops but are located nearby, which may be caused by some miss alignment of anchors. **(d)** Chromatin looping profiles at *Nanog* gene locus detected by Hi-TrAC and Micro-C. **(e)** Distribution of distance between loop anchors for unique loops detected by Hi-TrAC and Micro-C. **(f)** Distribution of the number of PETs for Micro-C overlapped and unique loops with Hi-TrAC. **(g)** The fractions of chromatin loop anchors detected by Hi-TrAC (upper panel) and Micro-C (lower panel), which are located to potential enhancers, promoters, or non-accessible regions. mESC ATAC-seq (GSM1830114^114^) peaks were used to define promoters (peaks within 2 kb of TSS) and enhancers (distance of peaks to TSS > 2kb). “Other” indicates the loop anchor has no overlaps with ATAC-seq peaks. **(h)** Aggregation analysis of ATAC-seq, DNase-seq, CTCF and H3K27me3 ChIP-seq signals on loop anchors defined as “Other” in panel **g** for Hi-TrAC (left panel) and Micro-C (right panel). These Hi-TrAC anchors are potential weak ATAC-seq peaks missed by peak-calling, meanwhile Micro-C anchors are weak CTCF binding sites.

**Extended Data Fig. 2.**
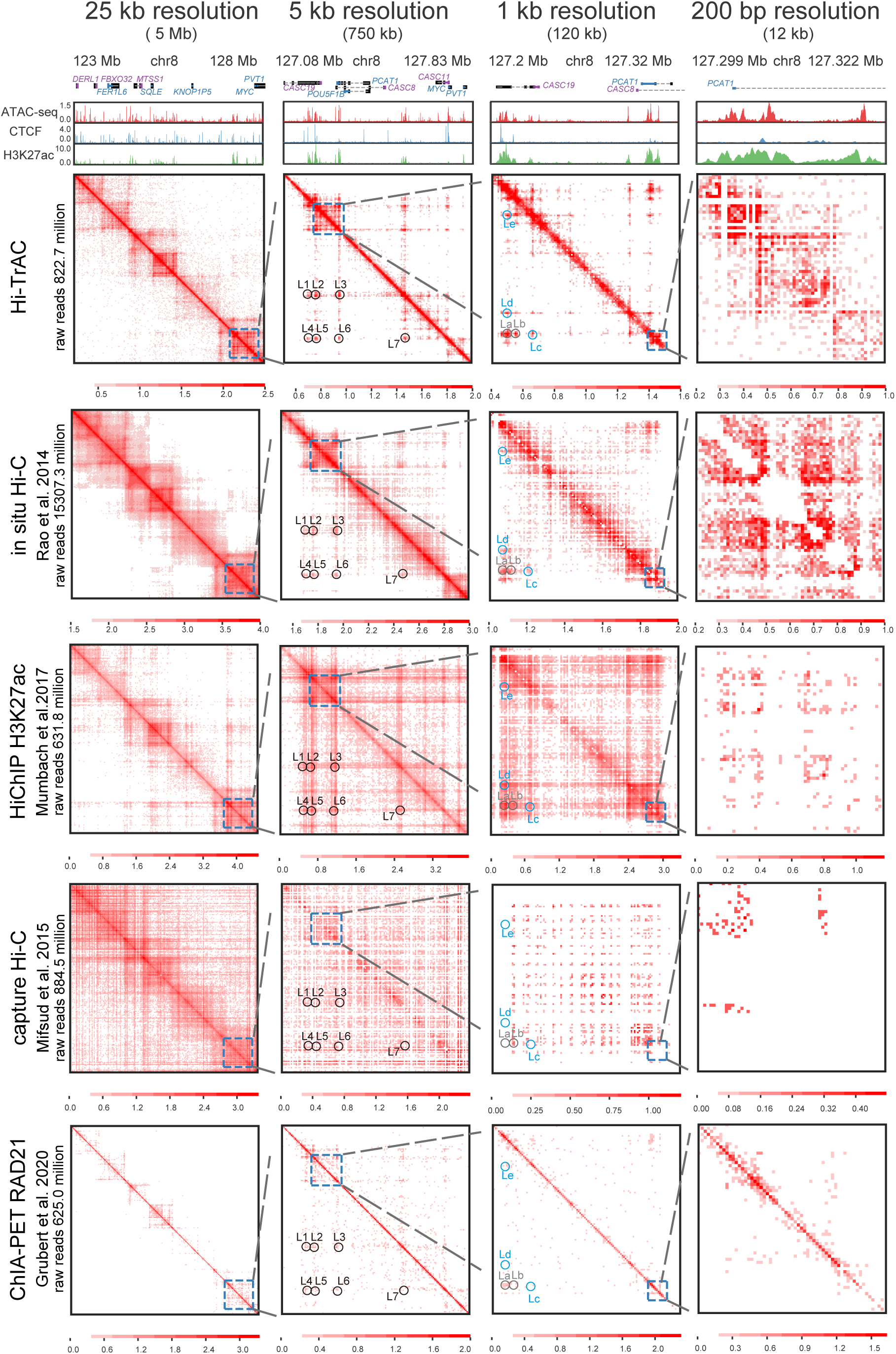
Comparison of Hi-TrAC to other techniques for mapping chromatin interactions. Comparison of chromatin architectures detected by Hi-TrAC and other representative state-of-art chromatin interactions mapping techniques, including in situ Hi-C^8^, H3K27ac HiChIP^65^, capture Hi-C^64^ and RAD21 ChIA-PET^63^, at different resolutions and scales in GM12878 cells around *MYC* gene locus. Numbers of raw sequenced reads were indicated for each method (refer to **Supplemental Table 1** for more details). Log_10_ transformed PETs were shown in the heatmaps. ChIP-seq data were obtained from the ENCODE project^115^. ATAC-seq data were obtained from GSE47753^109^. Visualization was performed with cLoops2 plot module.

**Extended Data Fig. 3.**
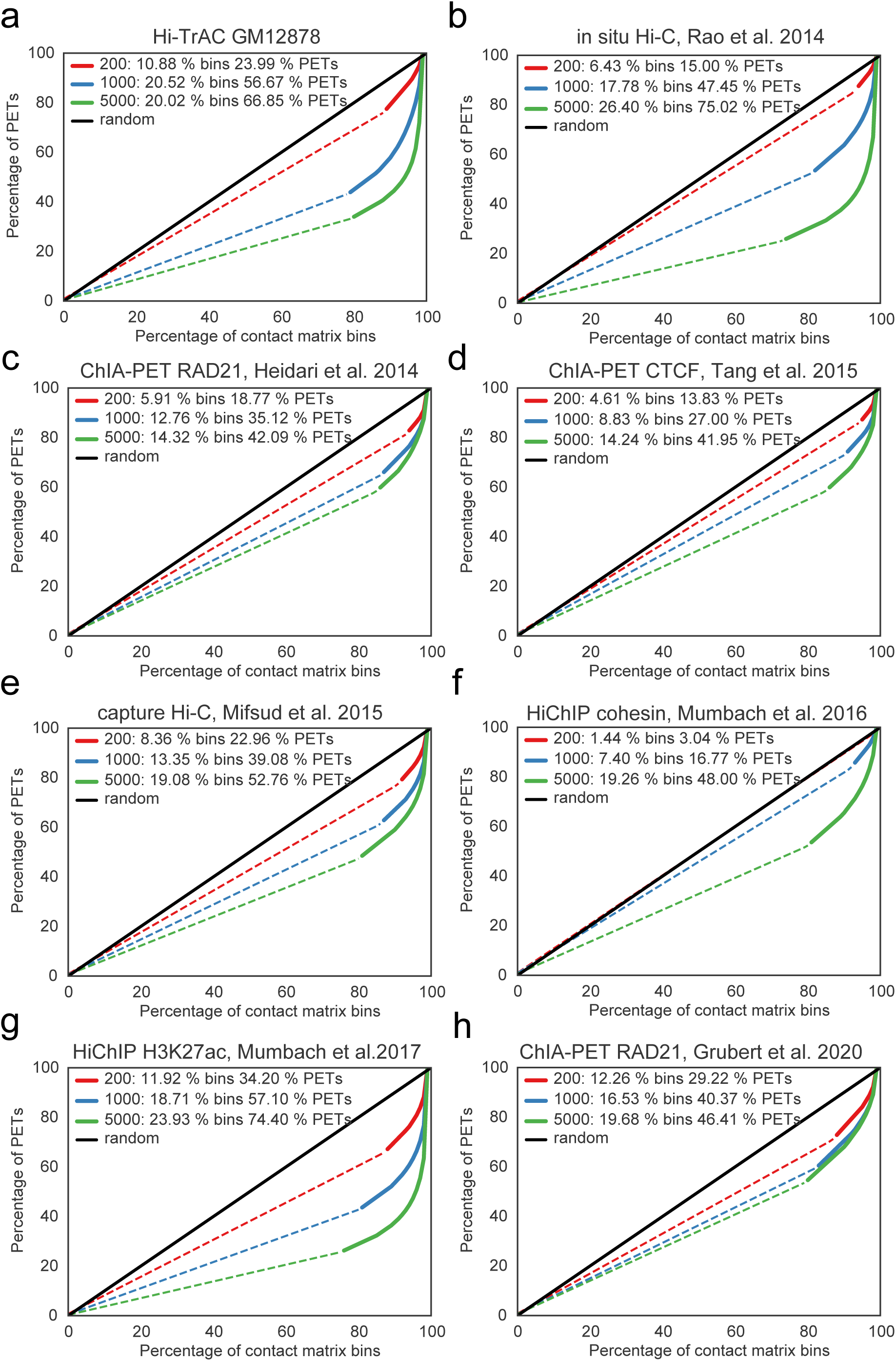
Estimation of resolutions of genome-wide interaction data from different techniques. Estimation of resolutions for Hi-TrAC (**a**), in situ Hi-C (**b**), RAD21 ChIA-PET (**c**), CTCF ChIA-PET (**d**), capture Hi-C (**e**), cohesin HiChIP (**f**), H3K27ac HiChIP (**g**) and RAD21 ChIA-PET (**h**). The interacting paired-end tags (PETs) were grouped into contact matrix bins based on different resolutions (200bp, 1kb, and 5kb). Dash lines show the contact matrix bins with only singleton PETs, which are evenly distributed and increased linearly and are presumably background noises. Solid curves indicate the bins with multiple PETs. We define the highest genome-wide resolution as more than 50% of PETs (solid curves) are in multiple PET bins. The analysis is implemented in the cLoops2 estRes module.

**Extended Data Fig. 4.**
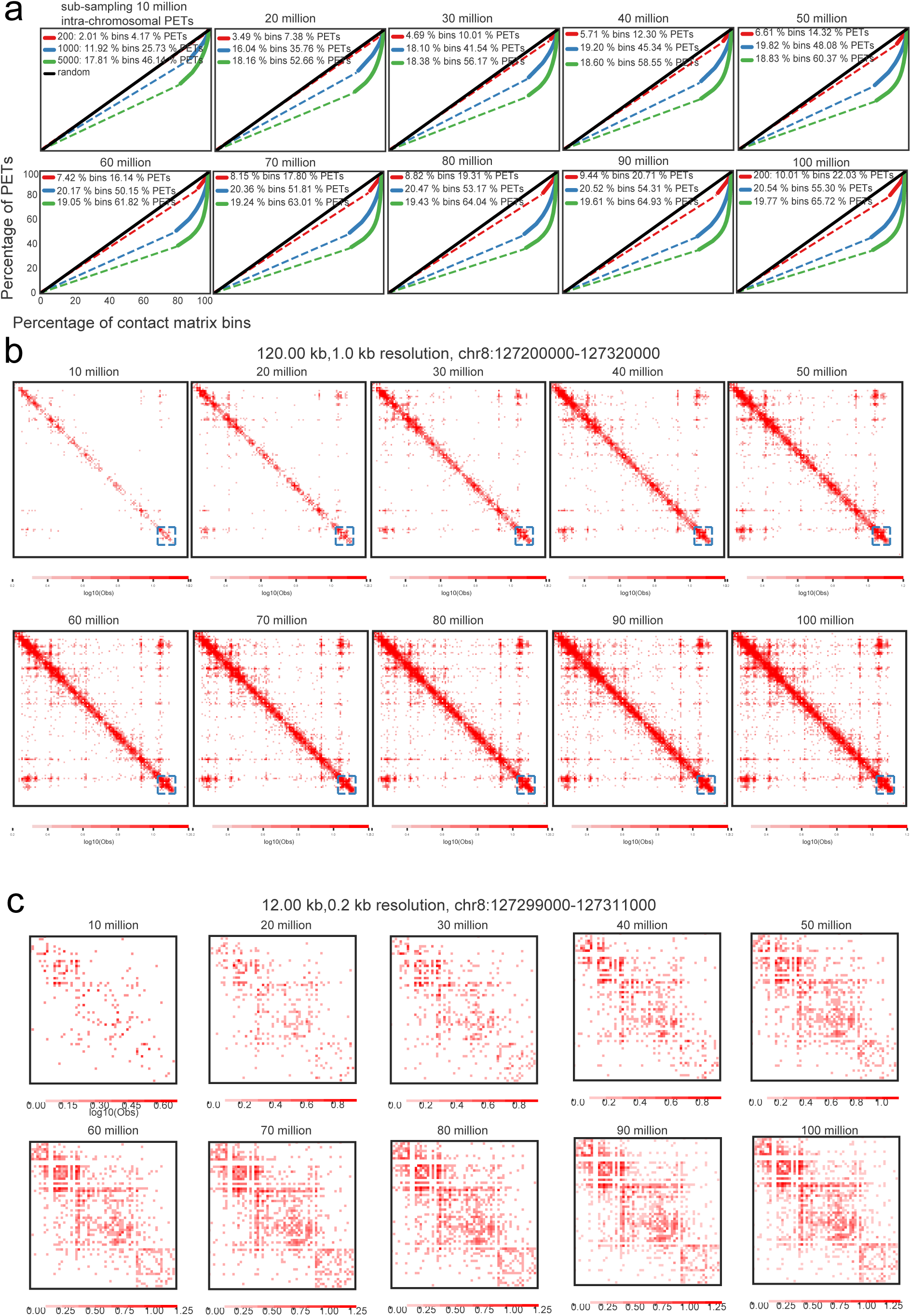
Downsampling of GM12878 Hi-TrAC data for estimating the required sequencing depth to achieve desired resolution. **(a)** Estimation of interaction resolutions with different sub-sampling depths of final unique intra-chromosomal PETs. With 60 million final unique intra-chromosomal PETs, Hi-TrAC can achieve the resolution of 1 kb. **(b)** The interaction matrix heatmaps for the example region shown in **Extended Data Fig.2** with 1 kb resolution at different sub-sampling depth. **(c)** The interaction matrix heatmaps for the super-enhancer region shown in **Extended Data Fig.2** at 200 bp resolution. With 50 million final unique intra-chromosomal PETs, Hi-TrAC can detect clear sub-structures of the super-enhancer.

**Extended Data Fig. 5.**
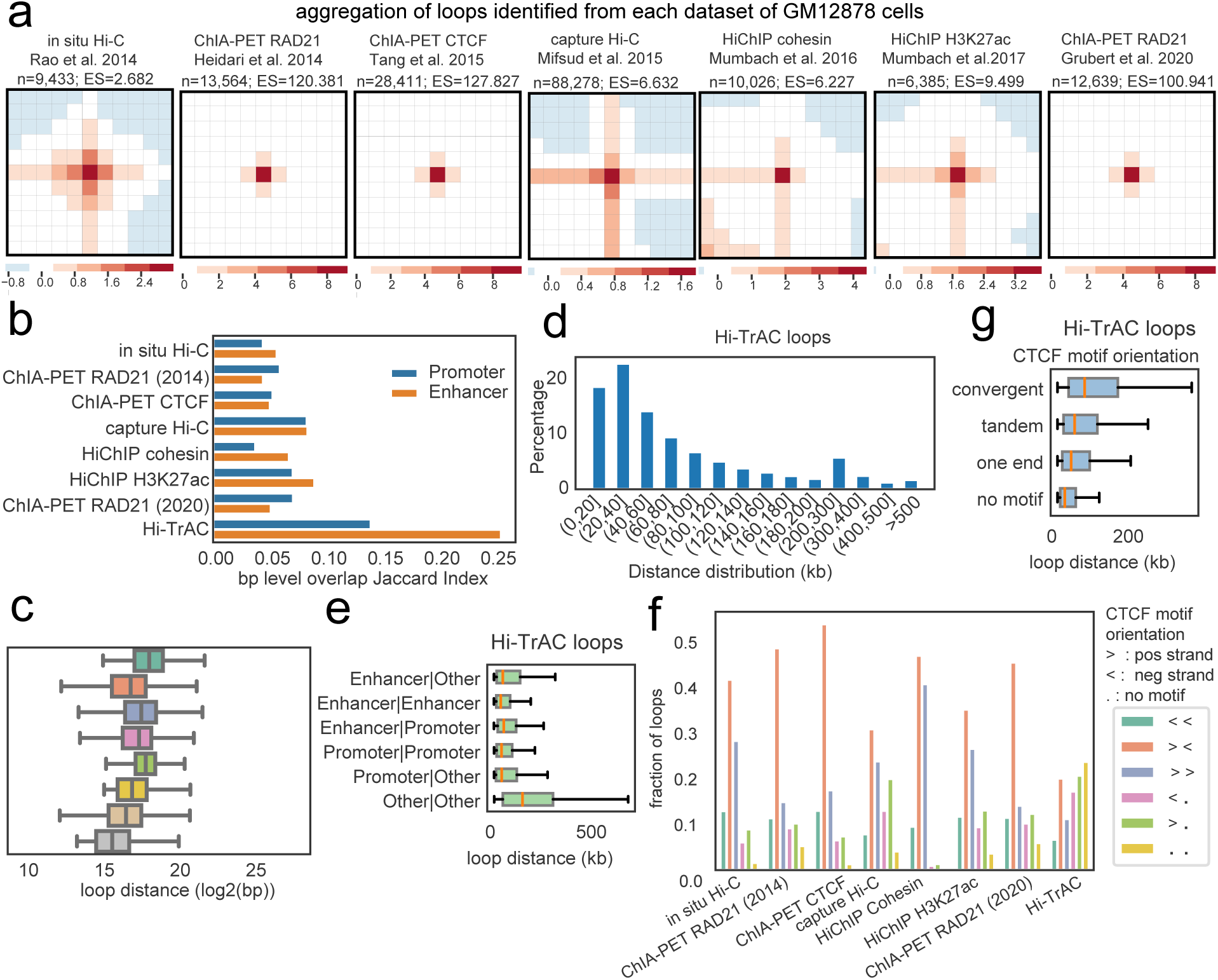
Features of chromatin loops detected by Hi-TrAC and other techniques. **(a)** Aggregation analysis of loops identified by other techniques. The numbers of loops called from each dataset are indicated. **(b)** Overlaps of loop anchors and cis-regulatory elements at base-pair level. A higher Jaccard index indicates anchor and enhancer/promoter matches with more similar size. **(c)** Distribution of anchor distance of loops detected by different techniques. **(d)** Distance distribution of GM12878 Hi-TrAC loop anchors. Most of the loops are formed by anchors within 200 kb. **(e)** Distance distribution of GM12878 loop anchors classified by cis-regulatory elements. **(f)** Summary of loop compositions with regard to CTCF motif orientation of two anchors. **(g)** Distance distribution of GM12878 loop anchors classified by CTCF motif orientation combinations.

**Extended Data Fig. 6.**
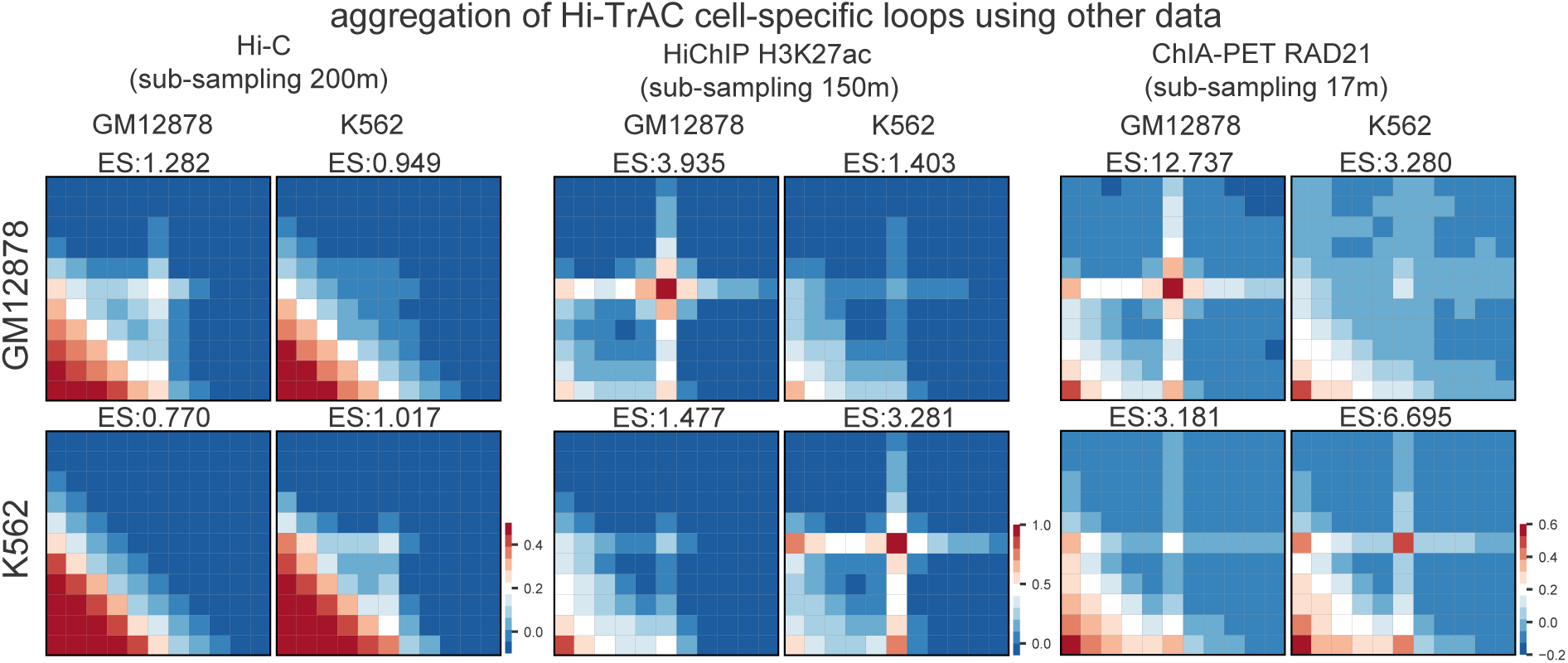
Validation of Hi-TrAC identified cell-specific loops. Aggregated loops analysis of Hi-TrAC cell-specific loops (**Supplemental Table 4**) using in situ Hi-C, HiChIP H3K27ac, and RAD21 ChIA-PET data. Enrichment score (ES) is calculated as the loop signal (the number of PETs at the matrix center) divided by nearby background (mean of the rest of the matrix except the center). ES > 1 indicates loops have relatively more interactions than nearby regions. A higher difference of ES indicates more differences of interacting PETs in the loops.

**Extended Data Fig. 7.**
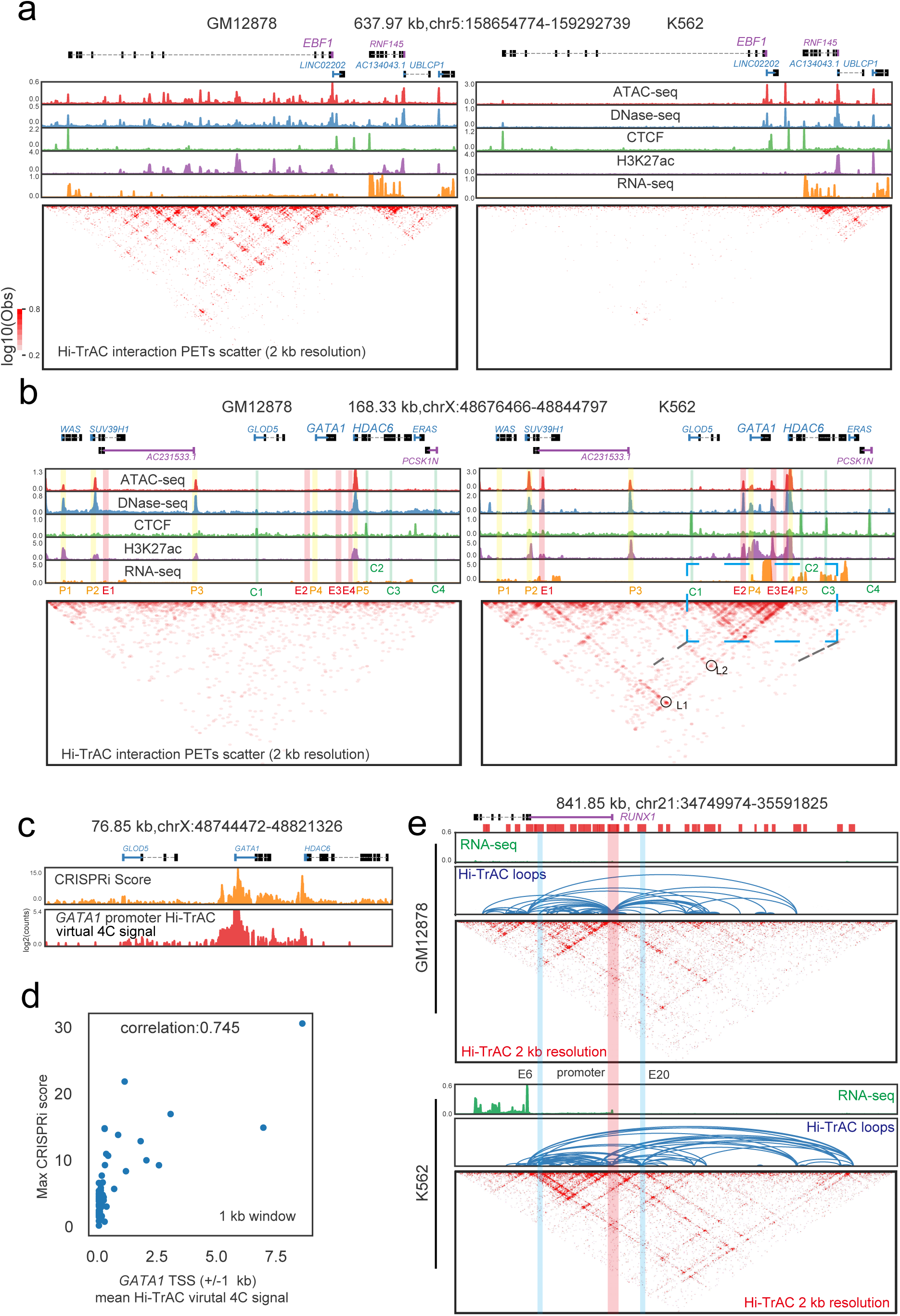
Hi-TrAC detects cell-specific interactions among regulatory elements at *EBF1*, *GATA1* and *RUNX1* gene loci. **(a)** The chromatin interaction profiles of *EBF1* gene locus detected by Hi-TrAC in GM12878 and K562 cells. ATAC-seq, DNase-seq, CTCF and H3K27ac ChIP-seq, and RNA-seq profiles are shown below the genomic annotations on the top, and interaction matrices are shown at the bottom. **(b)** The chromatin interaction profiles of *GATA1* gene locus detected by Hi-TrAC in GM12878 and K562 cells. Putative promoters were marked as P1 to P5, putative enhancers were marked as E1 to E4, and CTCF binding sites were marked as C1 to C4. The blue box region was highlighted for zoom-in presentation in panel **c**, and this region was studied before with CRISPR interference to identify regulatory elements of *GATA1* gene^67^. **(c)** Comparison of Hi-TrAC virtual 4C signals from *GATA1* promoter with CRISPRi scores reported previously^67^. **(d)** Correlation analysis of CRISPRi scores and Hi-TrAC virtual 4C signals from *GATA1* promoter. **(e)** Genome Browser snapshots showing looping profiles in GM12878 and K562 cells detected by Hi-TrAC at *RUNX1* gene locus.

**Extended Data Fig. 8.**
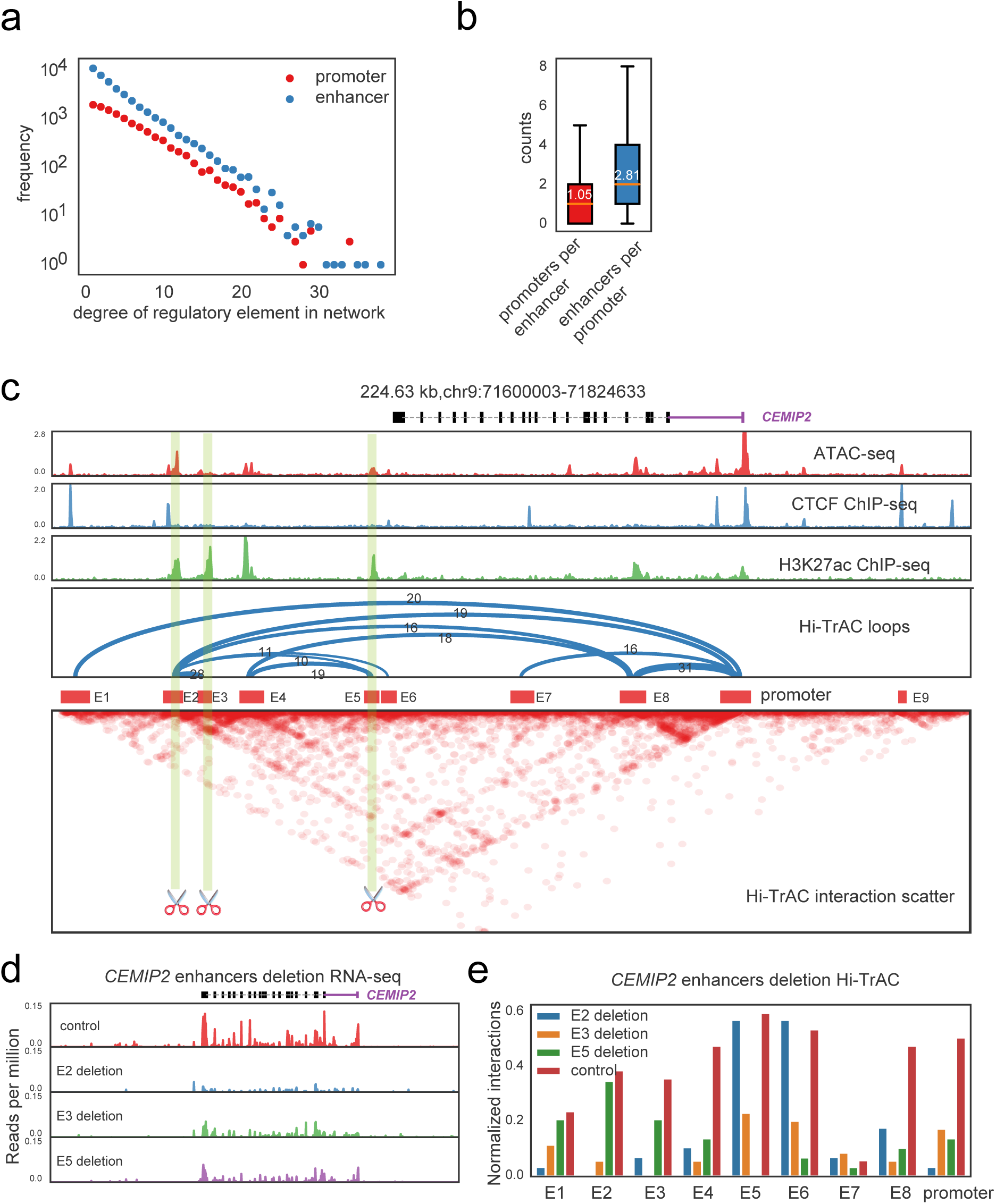
Regulation of gene expression by the promoter-enhancer interaction network. **(a)** Connection degree distributions of enhancers and promoters in the regulatory network constructed from GM12878 Hi-TrAC loops follow the scale-free network power-law. **(b)** Distributions of the numbers of looped targets per promoter or per enhancer. **(c)** Loops detected by Hi-TrAC for *CEMIP2* gene in K562 cells. Putative enhancers marked as E2, E3, and E5 were selected for deleting by CRISPR/Cas9. **(d)** RNA-seq assays showed decreased expression of *CEMIP2* by deleting E2, E3, or E5 as indicated in panel **c**. *ZNF234* promoter on a different chromosome was deleted as the negative control. **(e)** Interactions of *CEMIP2* promoter and enhancers decreased after the deletion of the putative enhancer E2, E3, or E5. The interactions were measured with Hi-TrAC data.

**Extended Data Fig. 9.**
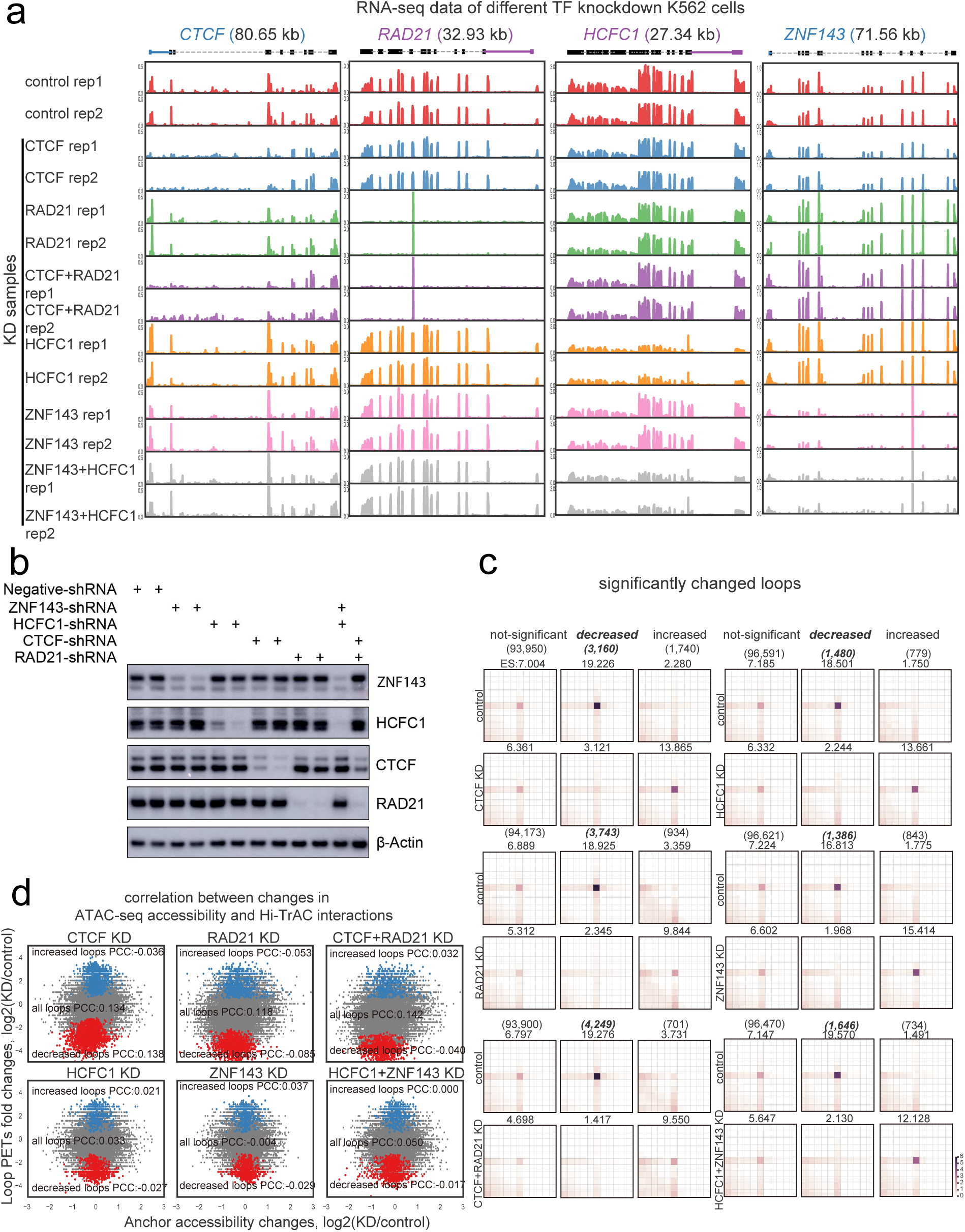
Knocking down ZNF143, HCFC1, CTCF, and RAD21 compromised chromatin looping in K562 cells. **(a)** K562 cells were infected with lentivirus carrying shRNA targeting either ZNF143, HCFC1, CTCF, or RAD21 alone or in combinations. The transcription of target genes were assessed by RNA-seq. **(b)** The expression of ZNF143, HCFC1, CTCF and RAD21 in knockdown cells were examined by western blotting. **(c)** Aggregation analysis of differentially enriched loops in knockdown cells (**Supplemental Table 6**). **(d)** Correlation analysis between the changes in accessibility measured by ATAC-seq and the changes in interactions measured by Hi-TrAC.

**Extended Data Fig. 10.**
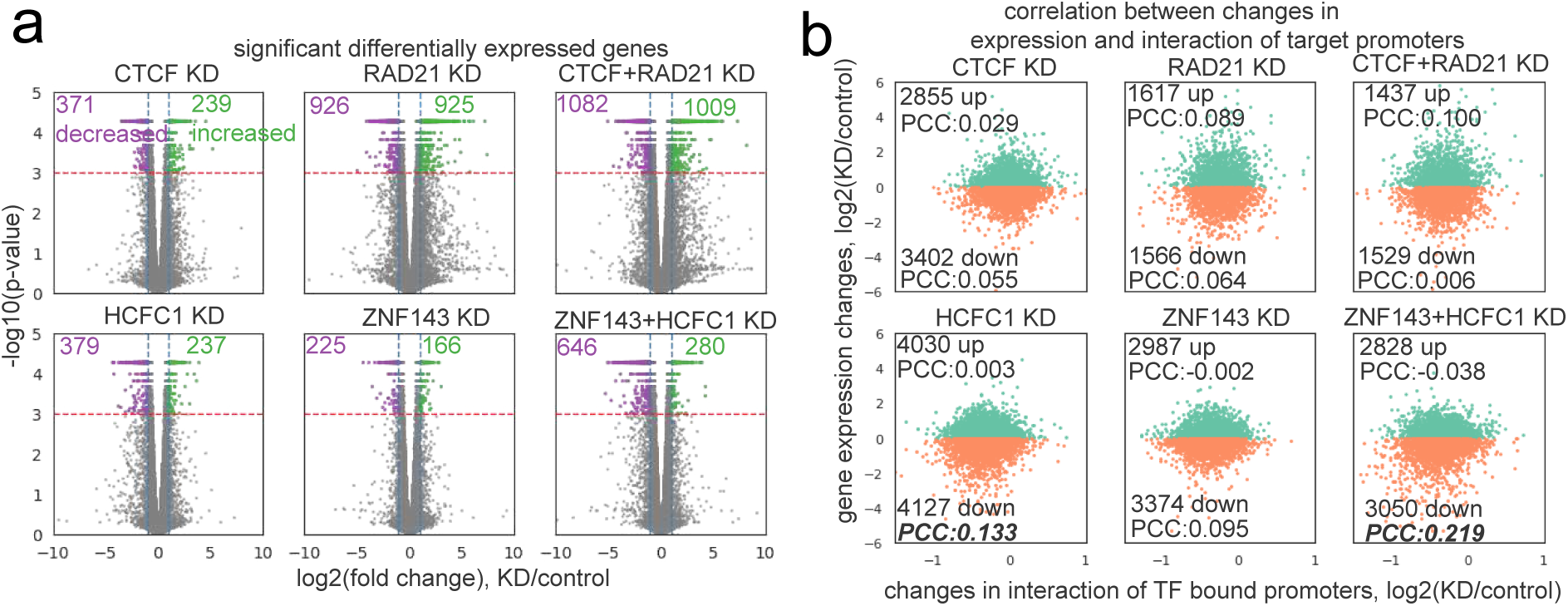
Gene expression impaired by knocking down CTCF, RAD21, HCFC1, and ZNF143. **(a)** Volcano plots showing significantly differentially expressed genes after knocking down CTCF, RAD21, HCFC1, and ZNF143 either alone or in combination. Purple dots indicate genes with decreased expression and green dots indicate genes with increased expression. **(b)** Correlation analysis between the changes in interactions at promoters and changes in gene expression. Interaction changes from Hi-TrAC data were measured for genes with promoters (+/-1Kb of TSS) overlapping with loop anchors bound by the targeted TF. PCC stands for Pearson Correlation Coefficient.

**Extended Data Fig. 11.**
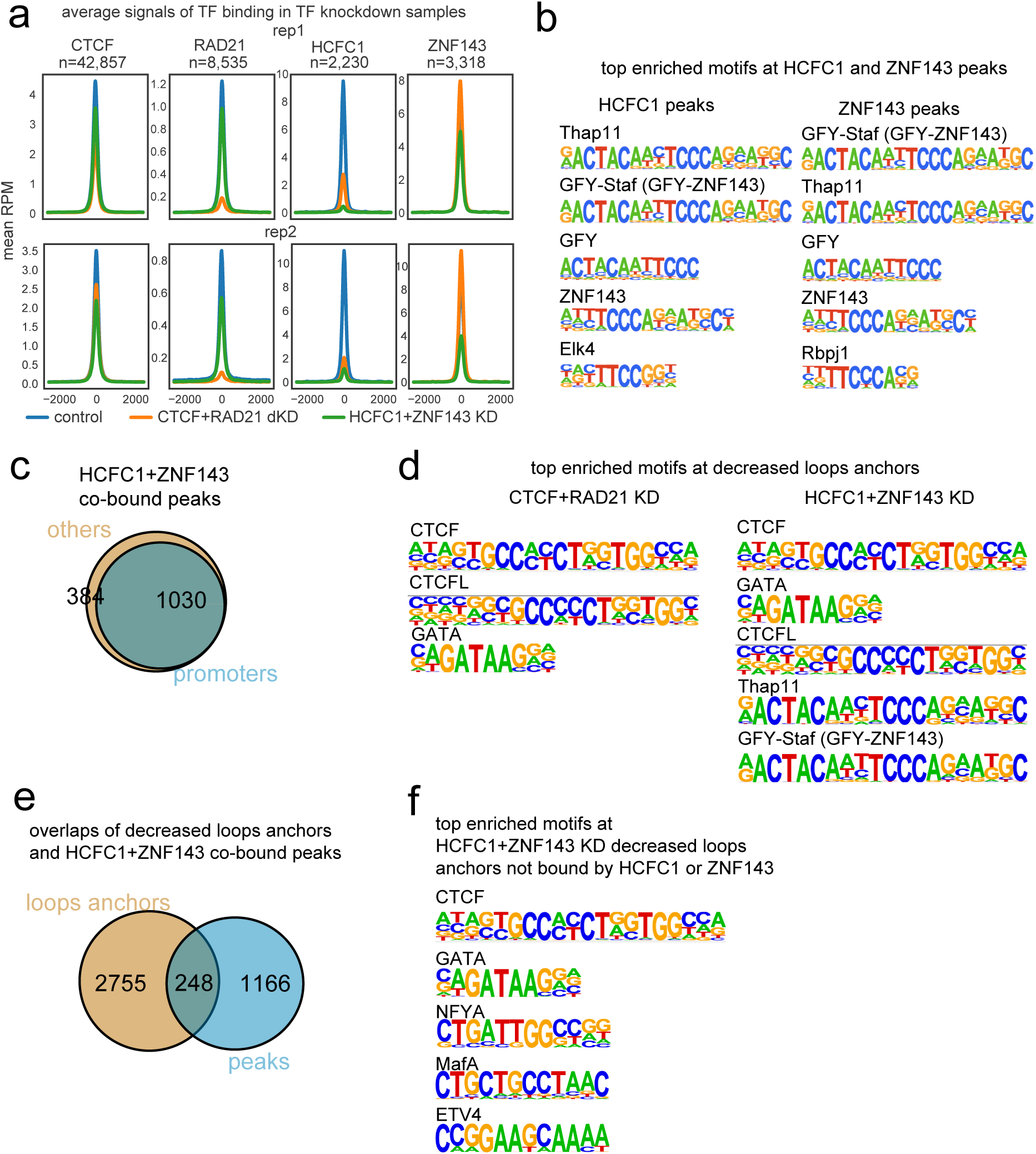
Characterization of loop anchors bound by different transcription factors. **(a)** Aggregated profile analysis for the changes of CTCF, RAD21, HCFC1 and ZNF143 binding after knocking down CTCF with RAD21, or HCFC1 with ZNF143. Overlapped peaks from control samples were used for the analysis. **(b)** Top enriched motifs found by HOMER from HCFC1 or ZNF143 peaks identified in K562 shRNA control cells. **(c)** Overlaps of HCFC1 and ZNF143 co-bound peaks and gene promoters. **(d)** Top enriched motifs found by HOMER from decreased loop anchors detected by Hi-TrAC in CTCF and RAD21 double knockdown (left panel) or HCFC1 and ZNF143 double knockdown (right panel) cells. **(e)** Overlaps of HCFC1 and ZNF143 co-bound peaks with significantly decreased loop anchors called from Hi-TrAC data after knocking down of HCFC1 and ZNF143. **(f)** Top enriched motifs in decreased loop anchors not bound by HCFC1 or ZNF143 in the HCFC1 and ZNF143 double knockdown cells.

**Extended Data Fig. 12.**
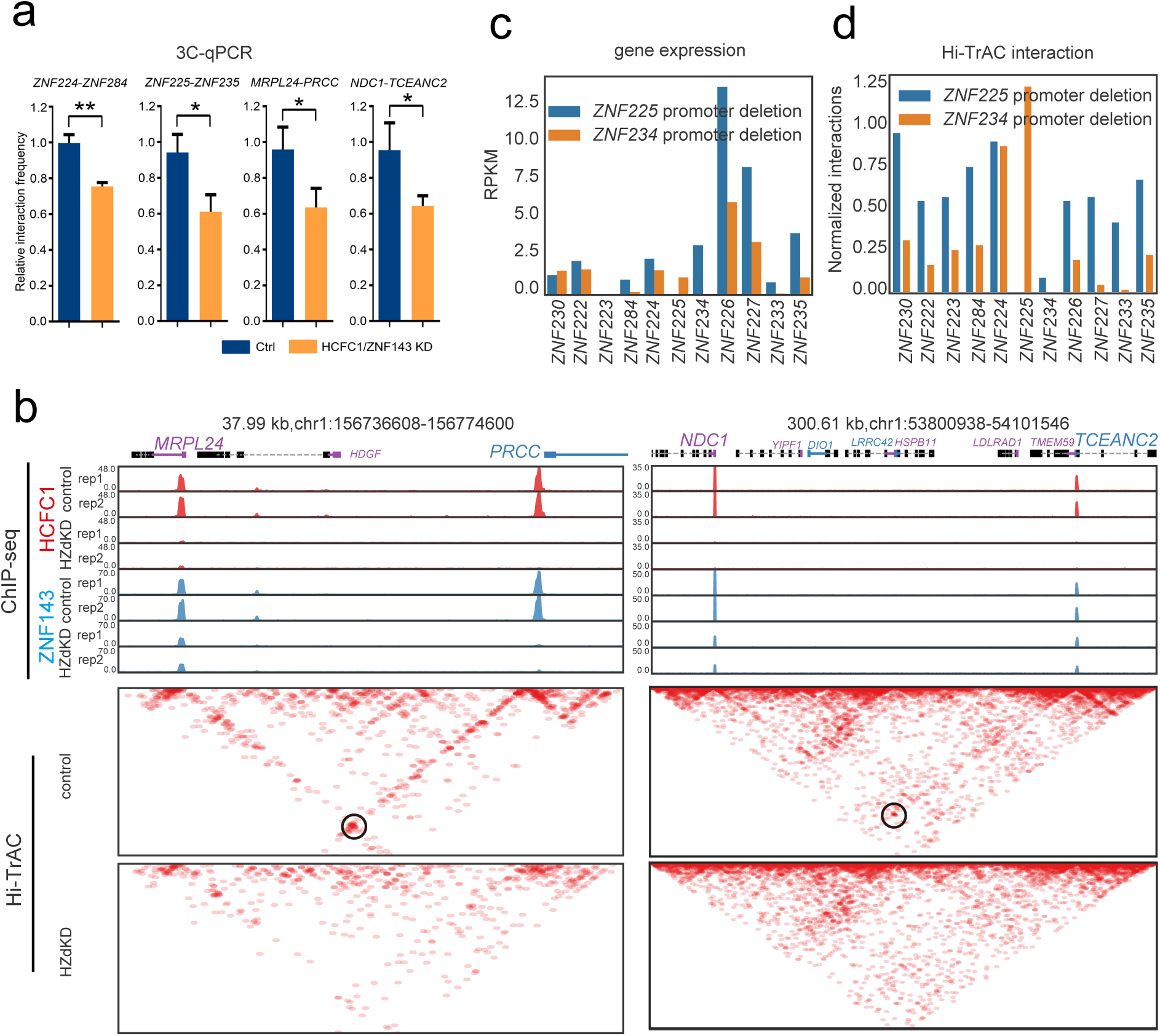
HCFC1 and ZNF143 associated promoter-promoter loops regulate gene expression. **(a)** 3C-qPCR quantitative analysis of the interaction frequency changes of *ZNF224*-*ZNF284*, *ZNF225*-*ZNF235*, *MRPL24*-*PRCC* and *NDC1*-*TCEANC2* in HCFC1 and ZNF143 knockdown cells. The abundance of these interaction pairs in 3C libraries were quantified by qPCR, and normalized to input control. The columns represent mean ± s.d., n = 3, ** p < 0.01, * p < 0.05 according to *t*-test. **(b)** Genome Browser snapshots showing *MRPL24*-*PRCC* and *NDC1*-*TCEANC2* loops impaired by knocking down HCFC1 and ZNF143 as detected by Hi-TrAC. The binding patterns of HCFC1 and ZNF143 (ChIP-seq tracks) at these regions are also presented. **(c)** The expression change of ZNF genes after deleting *ZNF225* or *ZNF234* promoter. **(d)** The chromatin interaction change of ZNF gene promoters after deleting *ZNF225* or *ZNF234* promoter.

## References

1 Naumova, N. et al. Organization of the mitotic chromosome. Science 342, 948–953, doi:10.1126/science.1236083 (2013).

2 Bonev, B. & Cavalli, G. Organization and function of the 3D genome. Nat Rev Genet 17, 661–678, doi:10.1038/nrg.2016.112 (2016).

3 Rowley, M. J. & Corces, V. G. Organizational principles of 3D genome architecture. Nat Rev Genet 19, 789–800, doi:10.1038/s41576-018-0060-8 (2018).

4 Gibcus, J. H. & Dekker, J. The hierarchy of the 3D genome. Mol Cell 49, 773–782, doi:10.1016/j.molcel.2013.02.011 (2013).

5 Misteli, T. The Self-Organizing Genome: Principles of Genome Architecture and Function. Cell 183, 28–45, doi:10.1016/j.cell.2020.09.014 (2020).

6 Zhang, H. et al. Chromatin structure dynamics during the mitosis-to-G1 phase transition. Nature 576, 158–162, doi:10.1038/s41586-019-1778-y (2019).

7 Lieberman-Aiden, E. et al. Comprehensive mapping of long-range interactions reveals folding principles of the human genome. Science 326, 289–293, doi:10.1126/science.1181369 (2009).

8 Rao, S. S. et al. A 3D map of the human genome at kilobase resolution reveals principles of chromatin looping. Cell 159, 1665–1680, doi:10.1016/j.cell.2014.11.021 (2014).

9 Rowley, M. J. et al. Evolutionarily Conserved Principles Predict 3D Chromatin Organization. Mol Cell 67, 837–852 e837, doi:10.1016/j.molcel.2017.07.022 (2017).

10 Sexton, T. et al. Three-dimensional folding and functional organization principles of the Drosophila genome. Cell 148, 458–472, doi:10.1016/j.cell.2012.01.010 (2012).

11 Nora, E. P. et al. Spatial partitioning of the regulatory landscape of the X-inactivation centre. Nature 485, 381–385, doi:10.1038/nature11049 (2012).

12 Dixon, J. R. et al. Topological domains in mammalian genomes identified by analysis of chromatin interactions. Nature 485, 376–380, doi:10.1038/nature11082 (2012).

13 Phillips-Cremins, J. E. et al. Architectural protein subclasses shape 3D organization of genomes during lineage commitment. Cell 153, 1281–1295, doi:10.1016/j.cell.2013.04.053 (2013).

14 Wijchers, P. J. et al. Cause and Consequence of Tethering a SubTAD to Different Nuclear Compartments. Mol Cell 61, 461–473, doi:10.1016/j.molcel.2016.01.001 (2016).

15 Parelho, V. et al. Cohesins functionally associate with CTCF on mammalian chromosome arms. Cell 132, 422–433, doi:10.1016/j.cell.2008.01.011 (2008).

16 Wendt, K. S. et al. Cohesin mediates transcriptional insulation by CCCTC-binding factor. Nature 451, 796–801, doi:10.1038/nature06634 (2008).

17 Zuin, J. et al. Cohesin and CTCF differentially affect chromatin architecture and gene expression in human cells. Proc Natl Acad Sci U S A 111, 996–1001, doi:10.1073/pnas.1317788111 (2014).

18 Sanborn, A. L. et al. Chromatin extrusion explains key features of loop and domain formation in wild-type and engineered genomes. Proc Natl Acad Sci U S A 112, E6456–6465, doi:10.1073/pnas.1518552112 (2015).

19 Fudenberg, G. et al. Formation of Chromosomal Domains by Loop Extrusion. Cell Rep 15, 2038–2049, doi:10.1016/j.celrep.2016.04.085 (2016).

20 Davidson, I. F. et al. DNA loop extrusion by human cohesin. Science 366, 1338–1345, doi:10.1126/science.aaz3418 (2019).

21 Ganji, M. et al. Real-time imaging of DNA loop extrusion by condensin. Science 360, 102–105, doi:10.1126/science.aar7831 (2018).

22 Vian, L. et al. The Energetics and Physiological Impact of Cohesin Extrusion. Cell 173, 1165–1178 e1120, doi:10.1016/j.cell.2018.03.072 (2018).

23 Kim, Y., Shi, Z., Zhang, H., Finkelstein, I. J. & Yu, H. Human cohesin compacts DNA by loop extrusion. Science 366, 1345–1349, doi:10.1126/science.aaz4475 (2019).

24 Zhang, Y. et al. The fundamental role of chromatin loop extrusion in physiological V(D)J recombination. Nature 573, 600–604, doi:10.1038/s41586-019-1547-y (2019).

25 Zhang, X. et al. Fundamental roles of chromatin loop extrusion in antibody class switching. Nature 575, 385–389, doi:10.1038/s41586-019-1723-0 (2019).

26 Dai, H. Q. et al. Loop extrusion mediates physiological Igh locus contraction for RAG scanning. Nature 590, 338–343, doi:10.1038/s41586-020-03121-7 (2021).

27 Fullwood, M. J. et al. An oestrogen-receptor-alpha-bound human chromatin interactome. Nature 462, 58–64, doi:10.1038/nature08497 (2009).

28 Handoko, L. et al. CTCF-mediated functional chromatin interactome in pluripotent cells. Nat Genet 43, 630–638, doi:10.1038/ng.857 (2011).

29 Chepelev, I., Wei, G., Wangsa, D., Tang, Q. & Zhao, K. Characterization of genome-wide enhancer-promoter interactions reveals co-expression of interacting genes and modes of higher order chromatin organization. Cell Res 22, 490–503, doi:10.1038/cr.2012.15 (2012).

30 DeMare, L. E. et al. The genomic landscape of cohesin-associated chromatin interactions. Genome Res 23, 1224–1234, doi:10.1101/gr.156570.113 (2013).

31 Dowen, J. M. et al. Control of cell identity genes occurs in insulated neighborhoods in mammalian chromosomes. Cell 159, 374–387, doi:10.1016/j.cell.2014.09.030 (2014).

32 Tang, Z. et al. CTCF-Mediated Human 3D Genome Architecture Reveals Chromatin Topology for Transcription. Cell 163, 1611–1627, doi:10.1016/j.cell.2015.11.024 (2015).

33 Javierre, B. M. et al. Lineage-Specific Genome Architecture Links Enhancers and Non-coding Disease Variants to Target Gene Promoters. Cell 167, 1369–1384 e1319, doi:10.1016/j.cell.2016.09.037 (2016).

34 Ji, X. et al. 3D Chromosome Regulatory Landscape of Human Pluripotent Cells. Cell Stem Cell 18, 262–275, doi:10.1016/j.stem.2015.11.007 (2016).

35 Smith, E. M., Lajoie, B. R., Jain, G. & Dekker, J. Invariant TAD Boundaries Constrain Cell-Type-Specific Looping Interactions between Promoters and Distal Elements around the CFTR Locus. Am J Hum Genet 98, 185–201, doi:10.1016/j.ajhg.2015.12.002 (2016).

36 Fukaya, T., Lim, B. & Levine, M. Enhancer Control of Transcriptional Bursting. Cell 166, 358–368, doi:10.1016/j.cell.2016.05.025 (2016).

37 Andrey, G. et al. Characterization of hundreds of regulatory landscapes in developing limbs reveals two regimes of chromatin folding. Genome Res 27, 223–233, doi:10.1101/gr.213066.116 (2017).

38 Furlong, E. E. M. & Levine, M. Developmental enhancers and chromosome topology. Science 361, 1341–1345, doi:10.1126/science.aau0320 (2018).

39 Kojic, A. et al. Distinct roles of cohesin-SA1 and cohesin-SA2 in 3D chromosome organization. Nat Struct Mol Biol 25, 496–504, doi:10.1038/s41594-018-0070-4 (2018).

40 Schoenfelder, S. & Fraser, P. Long-range enhancer-promoter contacts in gene expression control. Nat Rev Genet 20, 437–455, doi:10.1038/s41576-019-0128-0 (2019).

41 Kim, S. & Shendure, J. Mechanisms of Interplay between Transcription Factors and the 3D Genome. Mol Cell 76, 306–319, doi:10.1016/j.molcel.2019.08.010 (2019).

42 Benabdallah, N. S. et al. Decreased Enhancer-Promoter Proximity Accompanying Enhancer Activation. Mol Cell 76, 473–484 e477, doi:10.1016/j.molcel.2019.07.038 (2019).

43 Oh, S. et al. Enhancer release and retargeting activates disease-susceptibility genes. Nature, doi:10.1038/s41586-021-03577-1 (2021).

44 Robson, M. I., Ringel, A. R. & Mundlos, S. Regulatory Landscaping: How Enhancer-Promoter Communication Is Sculpted in 3D. Mol Cell 74, 1110–1122, doi:10.1016/j.molcel.2019.05.032 (2019).

45 van Arensbergen, J., van Steensel, B. & Bussemaker, H. J. In search of the determinants of enhancer-promoter interaction specificity. Trends Cell Biol 24, 695–702, doi:10.1016/j.tcb.2014.07.004 (2014).

46 Haberle, V. & Stark, A. Eukaryotic core promoters and the functional basis of transcription initiation. Nat Rev Mol Cell Biol 19, 621–637, doi:10.1038/s41580-018-0028-8 (2018).

47 Li, Y. et al. The structural basis for cohesin-CTCF-anchored loops. Nature 578, 472–476, doi:10.1038/s41586-019-1910-z (2020).

48 Nora, E. P. et al. Targeted Degradation of CTCF Decouples Local Insulation of Chromosome Domains from Genomic Compartmentalization. Cell 169, 930–944 e922, doi:10.1016/j.cell.2017.05.004 (2017).

49 Haarhuis, J. H. I. et al. The Cohesin Release Factor WAPL Restricts Chromatin Loop Extension. Cell 169, 693–707 e614, doi:10.1016/j.cell.2017.04.013 (2017).

50 Rao, S. S. P. et al. Cohesin Loss Eliminates All Loop Domains. Cell 171, 305–320 e324, doi:10.1016/j.cell.2017.09.026 (2017).

51 Heidari, N. et al. Genome-wide map of regulatory interactions in the human genome. Genome Res 24, 1905–1917, doi:10.1101/gr.176586.114 (2014).

52 Bailey, S. D. et al. ZNF143 provides sequence specificity to secure chromatin interactions at gene promoters. Nat Commun 2, 6186, doi:10.1038/ncomms7186 (2015).

53 Zhou, Q. et al. ZNF143 mediates CTCF-bound promoter-enhancer loops required for murine hematopoietic stem and progenitor cell function. Nat Commun 12, 43, doi:10.1038/s41467-020-20282-1 (2021).

54 Weintraub, A. S. et al. YY1 Is a Structural Regulator of Enhancer-Promoter Loops. Cell 171, 1573–1588 e1528, doi:10.1016/j.cell.2017.11.008 (2017).

55 Beagan, J. A. et al. YY1 and CTCF orchestrate a 3D chromatin looping switch during early neural lineage commitment. Genome Res 27, 1139–1152, doi:10.1101/gr.215160.116 (2017).

56 Xie, D. et al. Dynamic trans-acting factor colocalization in human cells. Cell 155, 713–724, doi:10.1016/j.cell.2013.09.043 (2013).

57 Pan, X. et al. YY1 controls Igkappa repertoire and B-cell development, and localizes with condensin on the Igkappa locus. EMBO J 32, 1168–1182, doi:10.1038/emboj.2013.66 (2013).

58 Li, L. et al. YY1 interacts with guanine quadruplexes to regulate DNA looping and gene expression. Nat Chem Biol 17, 161–168, doi:10.1038/s41589-020-00695-1 (2021).

59 Lai, B. et al. Trac-looping measures genome structure and chromatin accessibility. Nat Methods 15, 741–747, doi:10.1038/s41592-018-0107-y (2018).

60 Hsieh, T. S. et al. Resolving the 3D Landscape of Transcription-Linked Mammalian Chromatin Folding. Mol Cell 78, 539–553 e538, doi:10.1016/j.molcel.2020.03.002 (2020).

61 Krietenstein, N. et al. Ultrastructural Details of Mammalian Chromosome Architecture. Mol Cell 78, 554–565 e557, doi:10.1016/j.molcel.2020.03.003 (2020).

62 Hua, P. et al. Defining genome architecture at base-pair resolution. Nature, doi:10.1038/s41586-021-03639-4 (2021).

63 Grubert, F. et al. Landscape of cohesin-mediated chromatin loops in the human genome. Nature 583, 737–743, doi:10.1038/s41586-020-2151-x (2020).

64 Mifsud, B. et al. Mapping long-range promoter contacts in human cells with high-resolution capture Hi-C. Nat Genet 47, 598–606, doi:10.1038/ng.3286 (2015).

65 Mumbach, M. R. et al. Enhancer connectome in primary human cells identifies target genes of disease-associated DNA elements. Nat Genet 49, 1602–1612, doi:10.1038/ng.3963 (2017).

66 Mumbach, M. R. et al. HiChIP: efficient and sensitive analysis of protein-directed genome architecture. Nat Methods 13, 919–922, doi:10.1038/nmeth.3999 (2016).

67 Fulco, C. P. et al. Systematic mapping of functional enhancer-promoter connections with CRISPR interference. Science 354, 769–773, doi:10.1126/science.aag2445 (2016).

68 Huning, L. & Kunkel, G. R. The ubiquitous transcriptional protein ZNF143 activates a diversity of genes while assisting to organize chromatin structure. Gene 769, 145205, doi:10.1016/j.gene.2020.145205 (2021).

69 Ye, B. et al. ZNF143 in Chromatin Looping and Gene Regulation. Front Genet 11, 338, doi:10.3389/fgene.2020.00338 (2020).

70 Myslinski, E., Gerard, M. A., Krol, A. & Carbon, P. Transcription of the human cell cycle regulated BUB1B gene requires hStaf/ZNF143. Nucleic Acids Res 35, 3453–3464, doi:10.1093/nar/gkm239 (2007).

71 Halbig, K. M., Lekven, A. C. & Kunkel, G. R. The transcriptional activator ZNF143 is essential for normal development in zebrafish. BMC Mol Biol 13, 3, doi:10.1186/1471-2199-13-3 (2012).

72 Ngondo-Mbongo, R. P., Myslinski, E., Aster, J. C. & Carbon, P. Modulation of gene expression via overlapping binding sites exerted by ZNF143, Notch1 and THAP11. Nucleic Acids Res 41, 4000–4014, doi:10.1093/nar/gkt088 (2013).

73 Parker, J. B., Yin, H., Vinckevicius, A. & Chakravarti, D. Host cell factor-1 recruitment to E2F-bound and cell-cycle-control genes is mediated by THAP11 and ZNF143. Cell Rep 9, 967–982, doi:10.1016/j.celrep.2014.09.051 (2014).

74 Hancock, M. L. et al. Insulin Receptor Associates with Promoters Genome-wide and Regulates Gene Expression. Cell 177, 722–736 e722, doi:10.1016/j.cell.2019.02.030 (2019).

75 Chern, T. et al. Mutations in Hcfc1 and Ronin result in an inborn error of cobalamin metabolism and ribosomopathy. Nat Commun 13, 134, doi:10.1038/s41467-021-27759-7 (2022).

76 Izumi, H. et al. Role of ZNF143 in tumor growth through transcriptional regulation of DNA replication and cell-cycle-associated genes. Cancer Sci 101, 2538–2545, doi:10.1111/j.1349-7006.2010.01725.x (2010).

77 Kawatsu, Y. et al. The combination of strong expression of ZNF143 and high MIB-1 labelling index independently predicts shorter disease-specific survival in lung adenocarcinoma. Br J Cancer 110, 2583–2592, doi:10.1038/bjc.2014.202 (2014).

78 Verma, V., Paek, A. R., Choi, B. K., Hong, E. K. & You, H. J. Loss of zinc-finger protein 143 contributes to tumour progression by interleukin-8-CXCR axis in colon cancer. J Cell Mol Med 23, 4043–4053, doi:10.1111/jcmm.14290 (2019).

79 Zhang, L. et al. ZNF143-Mediated H3K9 Trimethylation Upregulates CDC6 by Activating MDIG in Hepatocellular Carcinoma. Cancer Res 80, 2599–2611, doi:10.1158/0008-5472.CAN-19-3226 (2020).

80 Myslinski, E., Gerard, M. A., Krol, A. & Carbon, P. A genome scale location analysis of human Staf/ZNF143-binding sites suggests a widespread role for human Staf/ZNF143 in mammalian promoters. J Biol Chem 281, 39953–39962, doi:10.1074/jbc.M608507200 (2006).

81 Michaud, J. et al. HCFC1 is a common component of active human CpG-island promoters and coincides with ZNF143, THAP11, YY1, and GABP transcription factor occupancy. Genome Res 23, 907–916, doi:10.1101/gr.150078.112 (2013).

82 Vinckevicius, A., Parker, J. B. & Chakravarti, D. Genomic Determinants of THAP11/ZNF143/HCFC1 Complex Recruitment to Chromatin. Mol Cell Biol 35, 4135–4146, doi:10.1128/MCB.00477-15 (2015).

83 Whalen, S., Truty, R. M. & Pollard, K. S. Enhancer-promoter interactions are encoded by complex genomic signatures on looping chromatin. Nat Genet 48, 488–496, doi:10.1038/ng.3539 (2016).

84 Mourad, R. & Cuvier, O. TAD-free analysis of architectural proteins and insulators. Nucleic Acids Res 46, e27, doi:10.1093/nar/gkx1246 (2018).

85 Wilber, A., Nienhuis, A. W. & Persons, D. A. Transcriptional regulation of fetal to adult hemoglobin switching: new therapeutic opportunities. Blood 117, 3945–3953, doi:10.1182/blood-2010-11-316893 (2011).

86 Sankaran, V. G. & Orkin, S. H. The switch from fetal to adult hemoglobin. Cold Spring Harb Perspect Med 3, a011643, doi:10.1101/cshperspect.a011643 (2013).

87 Wutz, G. et al. Topologically associating domains and chromatin loops depend on cohesin and are regulated by CTCF, WAPL, and PDS5 proteins. EMBO J 36, 3573–3599, doi:10.15252/embj.201798004 (2017).

88 Schwarzer, W. et al. Two independent modes of chromatin organization revealed by cohesin removal. Nature 551, 51–56, doi:10.1038/nature24281 (2017).

89 Kubo, N. et al. Promoter-proximal CTCF binding promotes distal enhancer-dependent gene activation. Nat Struct Mol Biol 28, 152–161, doi:10.1038/s41594-020-00539-5 (2021).

90 Li, G. et al. Extensive promoter-centered chromatin interactions provide a topological basis for transcription regulation. Cell 148, 84–98, doi:10.1016/j.cell.2011.12.014 (2012).

91 Wu, Y. et al. Promoter-anchored chromatin interactions predicted from genetic analysis of epigenomic data. Nat Commun 11, 2061, doi:10.1038/s41467-020-15587-0 (2020).

92 Eun, B., Sampley, M. L., Good, A. L., Gebert, C. M. & Pfeifer, K. Promoter cross-talk via a shared enhancer explains paternally biased expression of Nctc1 at the Igf2/H19/Nctc1 imprinted locus. Nucleic Acids Res 41, 817–826, doi:10.1093/nar/gks1182 (2013).

93 Picelli, S. et al. Full-length RNA-seq from single cells using Smart-seq2. Nat Protoc 9, 171–181, doi:10.1038/nprot.2014.006 (2014).

94 Frankish, A. et al. GENCODE reference annotation for the human and mouse genomes. Nucleic Acids Res 47, D766–D773, doi:10.1093/nar/gky955 (2019).

95 Khan, A. & Zhang, X. dbSUPER: a database of super-enhancers in mouse and human genome. Nucleic Acids Res 44, D164–171, doi:10.1093/nar/gkv1002 (2016).

96 Roadmap Epigenomics, C., et al. Integrative analysis of 111 reference human epigenomes. Nature 518, 317–330, doi:10.1038/nature14248 (2015).

97 Heinz, S. et al. Simple Combinations of Lineage-Determining Transcription Factors Prime cis-Regulatory Elements Required for Macrophage and B Cell Identities. Molecular Cell 38, 576–589, doi:10.1016/j.molcel.2010.05.004 (2010).

98 Langmead, B. & Salzberg, S. L. Fast gapped-read alignment with Bowtie 2. Nature Methods 9, 357–U354, doi:10.1038/Nmeth.1923 (2012).

99 Cao, Y., Liu, S., Ren, G., Tang, Q. & Zhao, K. cLoops2: a full-stack comprehensive analytical tool for chromatin interactions. Nucleic Acids Res 50, 57–71, doi:10.1093/nar/gkab1233 (2022).

100 Durand, N. C. et al. Juicer Provides a One-Click System for Analyzing Loop-Resolution Hi-C Experiments. Cell Syst 3, 95–98, doi:10.1016/j.cels.2016.07.002 (2016).

101 Cao, Y. et al. Accurate loop calling for 3D genomic data with cLoops. Bioinformatics 36, 666–675, doi:10.1093/bioinformatics/btz651 (2020).

102 Quinlan, A. R. & Hall, I. M. BEDTools: a flexible suite of utilities for comparing genomic features. Bioinformatics 26, 841–842, doi:10.1093/bioinformatics/btq033 (2010).

103 Grant, C. E., Bailey, T. L. & Noble, W. S. FIMO: scanning for occurrences of a given motif. Bioinformatics 27, 1017–1018, doi:10.1093/bioinformatics/btr064 (2011).

104 Weirauch, M. T. et al. Determination and inference of eukaryotic transcription factor sequence specificity. Cell 158, 1431–1443, doi:10.1016/j.cell.2014.08.009 (2014).

105 Cheneby, J., et al. ReMap 2020: a database of regulatory regions from an integrative analysis of Human and Arabidopsis DNA-binding sequencing experiments. Nucleic Acids Res 48, D180–D188, doi:10.1093/nar/gkz945 (2020).

106 Servant, N. et al. HiC-Pro: an optimized and flexible pipeline for Hi-C data processing. Genome Biol 16, 259, doi:10.1186/s13059-015-0831-x (2015).

107 Dobin, A. et al. STAR: ultrafast universal RNA-seq aligner. Bioinformatics 29, 15–21, doi:10.1093/bioinformatics/bts635 (2013).

108 Trapnell, C. et al. Differential gene and transcript expression analysis of RNA-seq experiments with TopHat and Cufflinks. Nat Protoc 7, 562–578, doi:10.1038/nprot.2012.016 (2012).

109 Buenrostro, J. D., Giresi, P. G., Zaba, L. C., Chang, H. Y. & Greenleaf, W. J. Transposition of native chromatin for fast and sensitive epigenomic profiling of open chromatin, DNA-binding proteins and nucleosome position. Nat Methods 10, 1213-1218, doi:10.1038/nmeth.2688 (2013).

110 Ramirez, F. et al. deepTools2: a next generation web server for deep-sequencing data analysis. Nucleic Acids Res 44, W160–165, doi:10.1093/nar/gkw257 (2016).

111 Hagberg, A., Swart, P. & S Chult, D. Exploring network structure, dynamics, and function using NetworkX. (Los Alamos National Lab.(LANL), Los Alamos, NM (United States), 2008).

112 Hunter, J. D. Matplotlib: A 2D graphics environment. Computing in science & engineering 9, 90–95 (2007).

113 Waskom, M. L. Seaborn: statistical data visualization. Journal of Open Source Software 6, 3021 (2021).

114 Di Stefano, B. et al. C/EBPalpha creates elite cells for iPSC reprogramming by upregulating Klf4 and increasing the levels of Lsd1 and Brd4. Nat Cell Biol 18, 371–381, doi:10.1038/ncb3326 (2016).

115 Consortium, E. P. An integrated encyclopedia of DNA elements in the human genome. Nature 489, 57–74, doi:10.1038/nature11247 (2012).

